# The highly rugged yet navigable regulatory landscape of the bacterial transcription factor TetR

**DOI:** 10.1101/2023.08.25.554764

**Authors:** Cauã Antunes Westmann, Leander Goldbach, Andreas Wagner

## Abstract

Transcription factor binding sites (*TFBSs*) are important sources of evolutionary innovations. Understanding how evolution navigates the sequence space of such sites can be achieved by mapping TFBS adaptive landscapes. In such a landscape, an individual location corresponds to a TFBS bound by a transcription factor. The elevation at that location corresponds to the strength of transcriptional regulation conveyed by the sequence. We developed an *in vivo* massively parallel reporter assay to map the landscape of bacterial TFBSs. We applied this assay to the TetR repressor, for which few TFBSs are known. We quantify the strength of transcriptional repression for 17,765 TFBSs and show that the resulting landscape is highly rugged, with 2,092 peaks. Only a few peaks convey stronger repression than the wild type. Non-additive (epistatic) interactions between mutations are frequent. Despite these hallmarks of ruggedness, most high peaks are evolutionarily accessible. They have large basins of attraction and are reached by around 20% of populations evolving on the landscape. Which high peak is reached during evolution is unpredictable and contingent on the mutational path taken. This first in-depth analysis of a prokaryotic gene regulator reveals a landscape that is navigable but much more rugged than the landscapes of eukaryotic regulators.

**Significance:** Understanding how evolution explores the vast space of genotypic possibilities is a fundamental question in evolutionary biology. The mapping of genotypes to quantitative traits (such as phenotypes and fitness) allows us to delineate adaptive landscapes and their topological properties, shedding light on how evolution can navigate such vast spaces. In this study, we focused on mapping a transcription factor binding site (TFBS) landscape to gene expression levels, as changes in gene expression patterns play a crucial role in biological innovation. We developed a massively parallel reporter assay and mapped the first comprehensive in vivo gene regulatory landscape for a bacterial transcriptional regulator, TetR. Surprisingly, this landscape is way more rugged than those observed in eukaryotic regulators. Despite its ruggedness, the landscape remains highly navigable through adaptive evolution. Our study presents the first high-resolution landscape for a bacterial TFBS, offering valuable insights into the evolution of TFBS in vivo. Moreover, it holds promise as a framework for discovering new genetic components for synthetic biological systems.

## INTRODUCTION

Mapping the relationship between genetic and phenotypic variation is crucial to understand the evolutionary process. Genotype-phenotype maps are widely used to study this relationship by connecting a potentially large genotype space to phenotypic traits, such as gene expression levels or enzymatic activity ^1,2^. If a trait is a scalar quantity, it can define an elevation at each coordinate (genotype). A special case of the resulting landscape is a fitness or adaptive landscape, in which the trait is an organism’s fitness ^3^. Evolution on such a landscape can be conceptualized as a hill-climbing process, in which populations are driven towards high fitness genotypes by natural selection ^4^. A smooth and single-peaked adaptive landscape facilitates the discovery of high fitness genotypes. Such a landscape is highly *navigable*^1^, because selection can lead a population to the global peak through multiple small “uphill” mutations. In contrast, a rugged landscape can hinder progress towards the highest peak, trapping populations on local suboptimal peaks with low fitness^1,5^.

New patterns of gene regulation are important sources of biological innovations ^1,6–8^. Transcriptional regulation is mainly controlled by transcription factors (TFs) that bind short DNA sequences known as transcription factor binding sites (TFBSs). Such binding helps regulate – activate or repress – gene expression ^9,10^. Therefore, mutational changes in TFBSs can play important roles in development, disease, and evolution of novel traits ^1,6–8^. However, the molecular origins and evolution of such elements are still poorly understood. While comprehensive in vitro data on the binding of eukaryotic TFs to their binding sites have helped us understand eukaryotic genotype-phenotype maps of gene regulation ^1,11–13^, the topography of such landscapes remains unexplored in prokaryotes.

To reduce this knowledge gap, we mapped the first comprehensive adaptive landscape of TFBS variants for bacteria. We chose the tetracycline repressor (TetR) as our model system, a well-studied TF with only two cognate binding sites, *tetO1* and *tetO2* ^14–16^. In the absence of the antibiotic tetracycline, TetR binds independently to *tetO1* and *tetO2* ^17–19^, repressing its own transcription and the expression of the *tetA* gene, which encodes an antibiotic efflux pump ^20^. We focus here on *tetO2*, because it affects the regulation of the *tet* genes more strongly ^17–19,21,22^. We refer to it as our wild-type binding site.

To characterize the TetR gene regulation landscape, we used a fluorescence-based *in vivo* method known as sort-seq ^23–25^ to map thousands of *tetO2* variants to gene expression levels. Because mutations that increase the affinity of a TF to its TFBS are often under positive selection, we assume that strong regulation and thus, strong binding are associated with high fitness ^26–30^. This assumption is further supported by the observation that strong repression of the *tetA* gene is essential to maintain high fitness in the absence of antibiotics ^31–33^.

We studied the topography of the TetR regulatory landscape and how accessible its peaks of strong repression are to adaptive evolution. We also simulated adaptive evolution on this landscape to determine whether evolving populations can easily find highly active binding sites. Our observations reveal that the landscape is highly rugged. Such rugged landscapes are thought to impede navigability ^1,34^. However, the highest peaks of this landscape – the most strongly repression tetR binding sites – are surprisingly accessible to adaptive evolution.

## RESULTS

### Experimental design

To map the regulatory landscape of TetR in vivo, we engineered a plasmid-based system for sort-seq experiments (**Figures 1a-b, Supplementary Figures S1-S2, Supplementary Methods 1-4**). This system allowed us to measure the binding of TetR and the resulting transcriptional repression for each genotype in a library of TetR TFBS variants, using fluorescence as a readout in a flow cytometry assay. The higher the affinity between TetR and its binding site, the lower the resulting GFP expression. To explore the TetR adaptive landscape, we randomized 8 symmetrically spaced base pair positions that are especially important for the binding of the *tetO2* sequence^15,21,35^. This resulted in a library size of 65,536 unique sequences, which constitutes the genotype space we analyzed (**Figure 1c, Supplementary Methods 5-6**).

**Figure 1.**
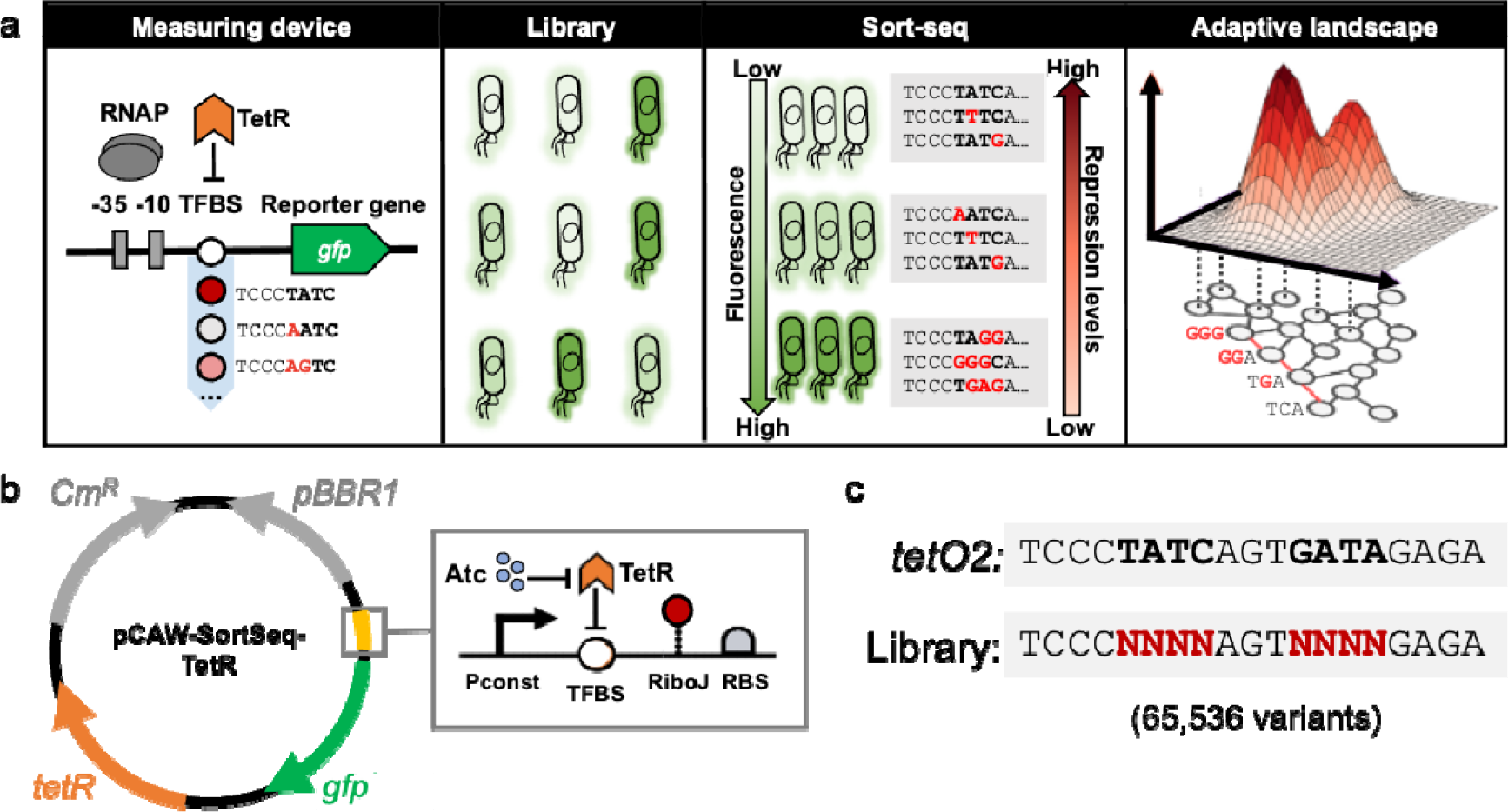
Experimental workflow. a. Sort-seq allows the mapping of gene regulatory landscapes. We designed a modular genetic system to quantify how strongly a prokaryotic transcriptional repressor can repress gene expression. In this system, both the repressor and its TFBSs can be easily replaced. The system is based on transcriptional repression by steric hindrance, because the cloned TFBS (or a library thereof) is positioned between the core promoter region (indicated as positions -35 and -10 from the transcription start site) that is recognized by the _s_70 factor of RNA polymerase (RNAP), and the *gfp* gene. Once the regulator binds to the TFBS, it physically blocks the activity of the RNA polymerase, reducing GFP expression. A library of thousands of TFBS variants results in a population of cells that differ in their GFP expression. We sorted this population in a fluorescence-activated cell-sorter (FACS) into 13 equidistant bins (on a binary logarithmic scale), where each bin harbors a subpopulation within a range of fluorescence intensities. We extracted, barcoded, and sequenced the DNA sequences (genotypes) from each bin. We mapped sequences to phenotypes (repression levels/binding scores) based on the sequence counts among different bins. We then built a network of genotypes (grey circles) in which sequences that differ in a single base pair are connected by an edge. The red edges in the network highlight a group of genotypes connected by single substitutions. This network, together with the repression level conveyed by each sequence constitutes the adaptive landscape we analyze. **b. The plasmid system**. The plasmid we developed for this study encodes a chloramphenicol resistance gene as a selective marker (Cm^R^, grey, right-oriented arrow), a broad-host low-copy replication origin (pBBR1, grey, left-oriented arrow), and two main modules interconnected by regulatory interactions. The first module contains the constitutively expressed *tetR* gene (orange). The TetR regulator expressed from this gene cassette acts on the second regulatory module of the system (yellow and green). In the regulatory region of this module (yellow, magnified in the grey box), TetR interacts with its cognate TFBS (or a TFBS variant from a library) and represses transcription (blunt vertical arrow). The addition of anhydrotetracycline (Atc, in blue) inhibits the repression from TetR (blunt horizontal arrow) and promotes transcription. Transcriptional effects from the TFBS sequences on the mRNA of *gfp* (green) are insulated by the RiboJ transcriptional insulator. The ribosome binding site (RBS) for the *gfp* gene is also shown. **c. Library overview**. We designed the *tetO2*-derived library by randomizing 8 symmetrically-spaced positions of the *tetO2* binding site (black bold letters) that are known to be important for TetR recognition^21,35^. The randomized sites (red bold Ns) are two symmetric palindromes of 4 base pairs each, yielding 4^8^= 65,536 library sequences.

### Sort-seq allows the high-resolution measuring of thousands of *tetO2* variants and the creation of PWMs

To assess our plasmid system’s ability to capture differences in repression levels of TetR binding site variants, we first measured the GFP expression driven by the wild-type sequence and four previously characterized variants cloned into our plasmid. These measurements showed expression levels consistent with previous reports ^58^ (**see Supplementary Figure S3**).

For subsequent experiments, we used our plasmid system with the wild-type binding site as a positive control for strong transcriptional repression and a negative control without a GFP promoter to set the lower bound of fluorescence as the basal autofluorescence of bacterial cells (**see Methods, Supplementary Methods 7 and Supplementary Figure S4**). We then proceeded to analyze the fluorescence of our library.

In the absence of the inducer, the wild type promotes strong repression (low fluorescence) (**Figure 2a**, left panel), whereas the entire TFBS library shows fluorescence that varies broadly (**Figure 2a**, right panel). This is the expected behavior if some variants lead to strong repression but others to only weak repression. In the presence of inducer, both the wild type and the entire library experience strong de-represssion (increased fluorescence), which reflects the expected dissociation of TetR from DNA.

**Figure 2.**
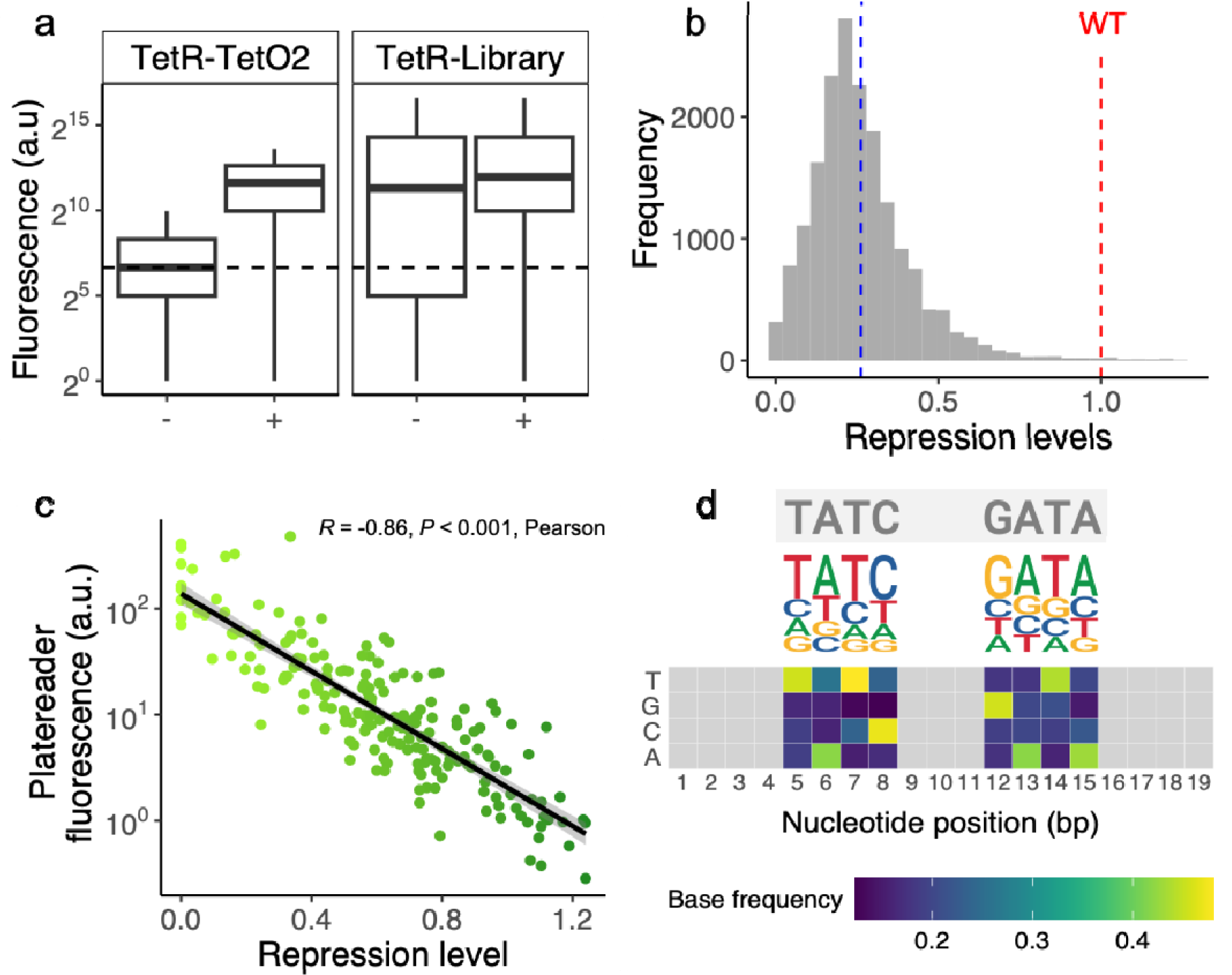
Our expression system captures repression by TetR binding sites. **a. Distribution of fluorescence levels between control and library**. Box plots summarize flow cytometry fluorescence measurements of the positive control (wild-type *tetO2*) and the library variants. Each box covers the range between the first and third quartiles (IQR). The horizontal line within the box represents the median value, and whiskers span 1.5 times the IQR. The dashed line represents the geometric mean (calculated for 3 replicates) of the autofluorescence of the negative control (promoterless GFP). We considered cells with values above this threshold as GFP expression-positive. The positive control (left panel, the *tetO2* binding site) strongly represses GFP expression in the absence of the inducer Atc (-). This repression is alleviated (fluorescence increases) in the presence of inducer (+). The same holds for the TFBS library (right panel), except that in the absence of the inducer, most library sequences bind the repressor more weakly than the wild type, resulting in higher fluorescence. **b. Distribution of repression levels (relative to *tetO2*) in the library**. The dashed blue line indicates the mean repression of all library variants (0.26), and the red dashed line indicates the repression level of the wild type. Data is based on a total of 17,851 variants. 78 variants (to the right of the red dashed line) repress expression more strongly than the wild type. **c. Plate reader validation of repression levels**. We selected a total of 195 mutants (15 mutants from each of 13 bins) and measured their fluorescence levels in a plate reader. We correlated their fluorescence measured in the plate reader (horizontal axis, arbitrary units, logarithmic scale) with our calculated repression level estimates (vertical axis, fold-change relative to wild type). *R* indicates the Pearson correlation coefficient (*R*^*2*^=-0.86). *P* indicates p-values for the correlation. The grey area around the regression line represents one standard error of the estimate at a confidence level of 95%. **d. Sequence logos and position frequency matrices for strongly repressing TetR binding sites**. Grey letters on top indicate the wild-type nucleotides for the library’s variable positions. We aligned the 78 sequences with repression levels higher than the wild type to construct a DNA sequence logo for strong TetR binding sites. The total height of the stack of letters depicts the information content of that nucleotide position, in bits. The proportions of all bases are shown as the relative heights of individual letters. The higher the relative height of a letter at a specific position, the more conserved and frequent the corresponding nucleotide is. Below the logo, we represent the corresponding frequency matrix for each variable position as a heat map. The vertical axis represents each base that can be found at each position (horizontal axis) of the TetR binding sites we examined. The heatmap color gradient represents the frequency of each base at each position (see color legend). Light grey cells represent binding site positions that were not mutated in the library.

We subdivided the cells from our library into 13 “bins”, based on their observed fluorescence values and sequenced variants from each bin (**see Methods, Supplementary Methods 7-8, Supplementary Figures S5-S6**). The resulting library contained 17,851 genotypes, which represent 27% of the 4^8^ genotypes in the genotype space we study (**see Methods**). Although we did not recover all genotypes, the correlation of read counts for each variant was remarkably consistent across replicates. The Pearson’s correlation coefficient R for replicates ranged from 0.971 to 0.991 (**Supplementary Figure S7**). We removed reads sequences with low read coverage (**see Methods, Supplementary Figures S8**). We used the observed distribution of individual variants among “bins” to map genotypes to their respective repression levels (**see Methods, Supplementary Methods 9, Supplementary Figures S9-S10**). We normalized repression levels by the wild type, such that values below one indicated weaker repression than the wild type, while values above one indicated stronger repression. To validate the data from our sort-seq experiments, we compared fluorescence levels from a plate reader assay with our sort-seq calculated repression levels for 15 variants from each bin, i.e., for a total of 13×15=195 variants. The correlation between the two measurements was strong and nearly linear (Pearson R=-0.86, p<0.001, N=195) (**Methods and Figure 2c**).

In our library, 78 variants showed stronger repression than the wild type, and we analyzed these variants to characterize the consensus of strong binding sites. We created frequency matrices for each nucleotide position based on an alignment of these variants. Displayed both as a heat map and sequence logo (**Figure 2d**), this frequency matrix reveals conserved nucleotides similar to the wild type (**Figure 2d**, grey) and additional nucleotides at each position that increased repression relative to the wild type. Our in vivo observations are consistent with previous in vitro data (**Supplementary Figure S11**).

### The TetR binding site adaptive landscape has multiple peaks

We used a network representation to study the adaptive landscape of the TetR binding sites (**Supplementary Methods 9.5**). In this representation, each node corresponds to a binding site variant with its respective level of repression. Variants separated by just one nucleotide (direct neighbors) are linked by a connection (edge) denoting a single mutation. Within this network, an evolutionary path consists of a chain of consecutive mutations. We found that the vast majority of the variants we examined (17,765, 99.5%) are part of the largest connected subgraph, also known as the “giant component”^51^. This particular subgraph represents our studied adaptive landscape (**Supplementary Table S1**).

A principal indicator of an adaptive landscape’s ruggedness is its number of peaks^1,36^. In our landscape, a peak is a genotype *g* whose neighbors all have a lower repression score than *g* itself (**see Supplementary Methods 9.5**). We found that the landscape has 2,092 peaks (12 percent of the total number of genotypes) and is thus highly rugged. The vast majority of these peaks (2,034/2,092, 97 percent) convey repression weaker than the wild type (**Figure 3a**). Only 58 peaks conveyed stronger repression than the wild type **(Figure 3a**). We refer to all 58 peaks with scores above *tetO2* as strong (repression) peaks or simply as high peaks.

**Figure 3.**
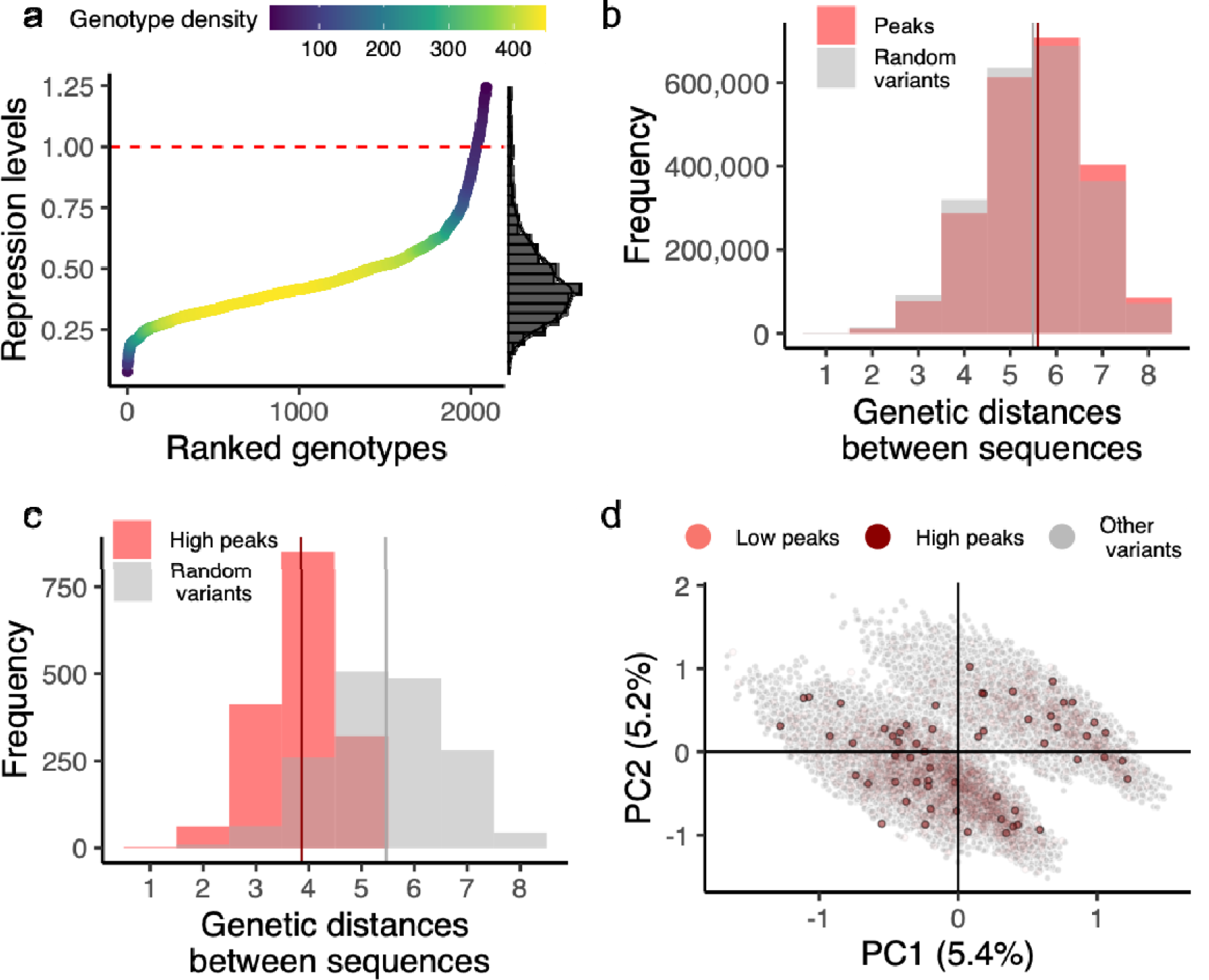
a. The TetR landscape has multiple adaptive peaks with widely varying repression strengths. Distribution of repression levels for 2,092 peaks in the landscape. The dashed red line at y=1 shows the repression level of the wild type. Gradient colors (see color legend) indicate the density of genotypes clustered along the horizontal axis. The histogram on the right represents the distribution of repression levels. **b. Peaks exhibit genetic variability**. The comparative genetic distances between nucleotide sequences are shown for 2,092 peaks (depicted in red) and for 2,092 random non-peak variants (in grey). The vertical lines represent the mean values for these distributions (d=5.59 for peaks [in red] against d=5.5 for the random variants [in grey]). **c. High repression peaks exhibit lesser genetic diversity than other variants**. The genetic spread is compared between nucleotide sequences of 58 high peaks (colored in red) and 58 randomly selected variants that exclude peaks (shown in grey). The mean values for these distributions are indicated by vertical lines (d=3.86 for the dominant peaks [in red] against d=5.46 for the random variants [in grey]). **d. A principal component analysis reveals that adaptive peaks are distributed widely within the genotype space** (**Supplementary Methods 9.6**). The panel shows principal components (PC) 1 and 2 and the amount of variation (in percentages) explained by each component. Every circle corresponds to one among the 17,765 variants within the landscape. Dark red and light red circles represent high and low peaks, respectively, while grey circles represent the other variants.

We investigated how the spatial arrangement of peaks in the landscape impacts their evolutionary accessibility. For this purpose, we examined if various peaks were closely positioned within the landscape, quantifying their genetic distances in pairwise comparisons. This metric signifies the least number of mutations needed to change one peak into another. For comparison, we also computed the distance distribution among pairs of 2,092 non-peak variants selected at random from the landscape. The mean distances were almost identical (*d*=5.59 for peaks vs. *d*=5.5 for random variants, (*d*=5.59 for peaks vs. *d*=5.5 for random variants, two-sided Kolmogorov–Smirnov test D = 0.03, p < 2.2 × 10^−16^, N_1_= 2,187,186, N_2_=2,187,186, **Figure 3b**). This pattern of similarity, however, did not apply to the high repression peaks. They are closer together in sequence space than expected by chance (*d*=3.86 for high peaks vs. *d*=5.46 for random variants, two-sided Mann–Whitney *U =* 412,662, p *<* 2.2 × 10^−16^, N_1_= 1,653, N_2_= 1,653, **Figure 3c**). However, their average distance of almost four mutational steps implies that high peaks are not confined to a small region of genotype space.

This can also be observed in **Figure 3d**, which displays the result of a principal component analysis (**Supplementary Methods 9.6**) showing how adaptive peaks are spread across the genotype space. The PCA reveals two “clouds” of genotypes **(Figure 3d)**, whose members are distinguished by the nucleotide at position 12 (**Figure 2d**). Genotypes with a G at position 12 fall into one cloud, whereas genotypes with A, C, or T at this position largely fall into the other cloud (**Supplementary Figure S12**). In sum, the TetR binding site landscape features numerous adaptive peaks spread out extensively, with high repression peaks being genetically more similar than other variants.

### High peaks are moderately accessible

A large population evolving by mutation and natural selection cannot traverse the valleys that lie between the high peaks of a rugged adaptive landscape, because it will get stuck along evolutionarily inaccessible evolutionary paths to a high peak, i.e., paths on which fitness decreases at least for some mutational steps. Such paths are inaccessible, at least in large populations like that of *E*.*coli* (with effective population size N=10^8^) ^37^, where even weakly deleterious mutations are unlikely to go to fixation^52^. Because the TetR landscape we study is rugged, the gradual evolution of strong TetR binding sites from weak ones might face this obstacle.

To find out whether this is the case, we studied the evolutionary accessibility of high peaks. Initially, we assessed if there were accessible paths from each variant to each high repression peak. Each mutational step along such a path would increase the repression strength of the evolving TetR binding site. We measured the size of each peak’s basin of attraction, indicating the number of variants through which the peak is reachable. (**Figure 4a**). We found that the basin sizes of high peaks vary widely, ranging from high peaks accessible from only a single variant (1/15,673) to others being accessible from 52.5% (8231/15,673) of all the variants **(Supplementary Figure S13**). Basin sizes varied considerably with peak repression strength (**Figure 4b**) and comprised, on average, 4.5 ± 6 (mean ±s.d.) percent of variants (946/15,673). Remarkably, low peaks generally had basins with a smaller size (median: 276, 1.7 % of all variants) than high peaks (median: 1260, 8%) (two-sided Mann– Whitney *U* = 90680, p = 2.81 × 10^−12^, N_1_=, 58 N_2_= 2,034) (**Figure 4b**). Moreover, when considering all peaks, we observed a moderate statistical association between basin size and the repression levels of peaks (Pearson *R*=0.43, *p* <0.001, N=2,092; **Figure 4c**).

**Figure 4.**
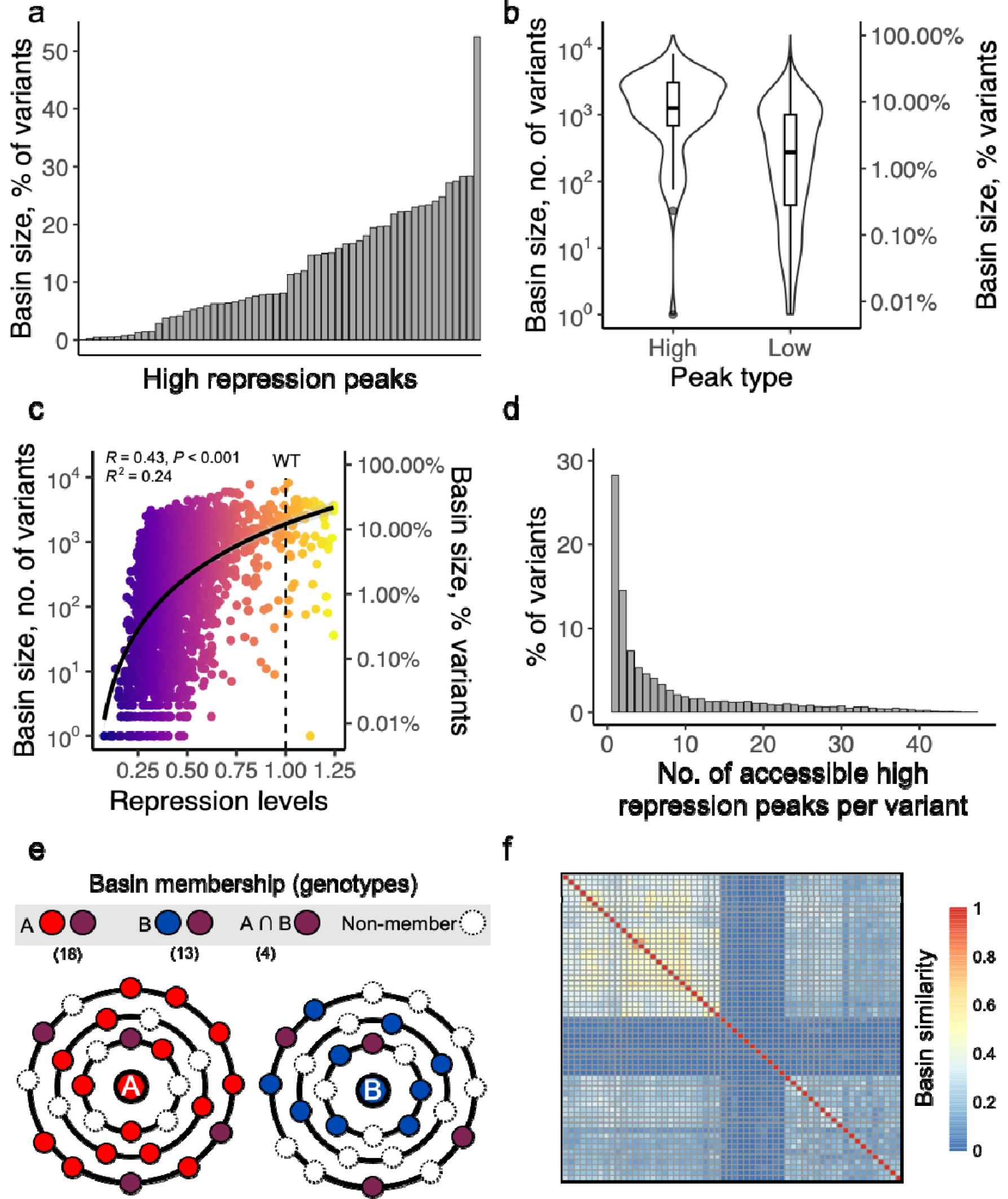
Basins of attraction. a. The basin sizes of high peaks. The basin size (vertical axis) of each peak (horizontal axis) is expressed as the percentage of TFBS variants that can access the peak. The peaks have been arranged along the x-axis in ascending order of their basin sizes. **b. High peaks have larger basin sizes than low peaks**. The left and right vertical axes show the basin sizes of peak genotypes as the number of variants (left) and the proportion of all variants (right). The horizontal axis subdivides the data into high peaks (N=58, repression stronger than *tetO2*) and low peaks (N=2,034, repression weaker than *tetO2*). The violin plots show the distribution of basin sizes for each of the two kinds of peaks. Their width represents the frequency of values. The vertical length of the box in each box covers the range between the first and third quartiles (IQR). The horizontal line within the box represents the median value, and whiskers span 1.5 times the IQR. **c. High peaks tend to have larger basins of attraction**. The scatter plot represents the association between repression levels conveyed by a peak variant (horizontal axis) and the size of its basin of attraction (absolute numbers on the left vertical axis, percentages on the right vertical axis, note logarithmic scale). The heatmap colors represent repression levels conveyed by each peak genotype from low (dark blue) to high (yellow). The dashed vertical line represents the repression level for the wild type. The black curve represents the linear regression model for the data, and the grey shade around it represents its 95 percent confidence interval. *R* is the Pearson correlation coefficient (*R*^*2*^ =0.24, N=2,092). **d. Many variants can attain more than a single peak**. We delineated the distribution of accessed high peaks per variant by quantifying the number of peaks potentially accessed by each variant **e. Schematic of basin size and overlaps between basins of attraction**. We computed the basins of attraction of each peak by counting the number of genotypes (sequence variants) from which an accessible path exists to each peak. We computed the overlap between the basins of attraction of two peaks with the Jaccard index (Supplementary Methods 9.5). Each circle represents a single hypothetical genotype. A (red) and B (blue) represent two hypothetical peaks. The concentric rings around A and B represent genotypes at increasing mutational distances from the peaks. Small red, blue, purple, and white circles (genotypes) indicate members of the basin of attraction of only peak A, only peak B, both peaks, or none of the peaks, respectively. The total number of genotypes for peak A, peak B and their intersection is indicated in parentheses **f. The attraction basins associated with several high peaks demonstrate significant overlap**. The heatmap matrix represents the fraction of TFBS variants concurrently found within the basins of attraction across all pairwise comparisons between the 58 high peaks (Methods). Each row and column correspond to one peak, and each matrix entry indicates the overlap (Jaccard index, Supplementary Methods 9.5) between the corresponding pairs of peaks. A value of one (represented in red) indicates that the attraction basins of the compared peaks contain the same variants, whereas a value of zero (portrayed in yellow) underscores that the two basins possess no mutual variants.

Given that high-repression peaks possess large basins of attraction, it’s expected for individual variants to belong to multiple basins, enabling them to reach several repression peaks. Specifically, this holds for 57.2% (8,965/15,673) of variants (**Figure 4d**).

To evaluate the intersection among distinct peak basins, we quantified how many variants are shared between them in a pairwise manner (**Figure 4e**). Such basin overlap varies widely, with some pairs of high peaks sharing 80% of the variants in their basins and others not sharing any variants. On average, the basins of high peaks shared 17%±14% (mean ±s.d.) of the variants in their basins **(Figure 4f)**. Although the sharing of basin members varied between peaks, the basins of high peaks shared more variants than low peaks (two-sided Mann–Whitney *U*□= 2,633,301,522, p < 2.2 × 10^−16^, N_1_=1,653, N_2_=2,067,561). In sum, high peaks are more accessible than low peaks and share a greater proportion of their basins of attraction.

In an ideally smooth landscape, the genetic distance between a variant and a peak equates to the shortest navigable route. However, within a multipeaked landscape like ours, feasible routes might be way longer than the genetic distance. In the regulatory landscape of TetR, the shortest accessible paths that terminate at a given high peak from any one variant are on average two mutations longer (mean±s.d.= 7.53±2.63 mutations) than the mean genetic distance between the variant and the peak (5.43±1.33 mutations) (**Figure 5a**, Welch Two Sample t-test, t = -289.92, df = 247078, p-value < 2.2 × 10^−16^, N= 166,417).

**Figure 5.**
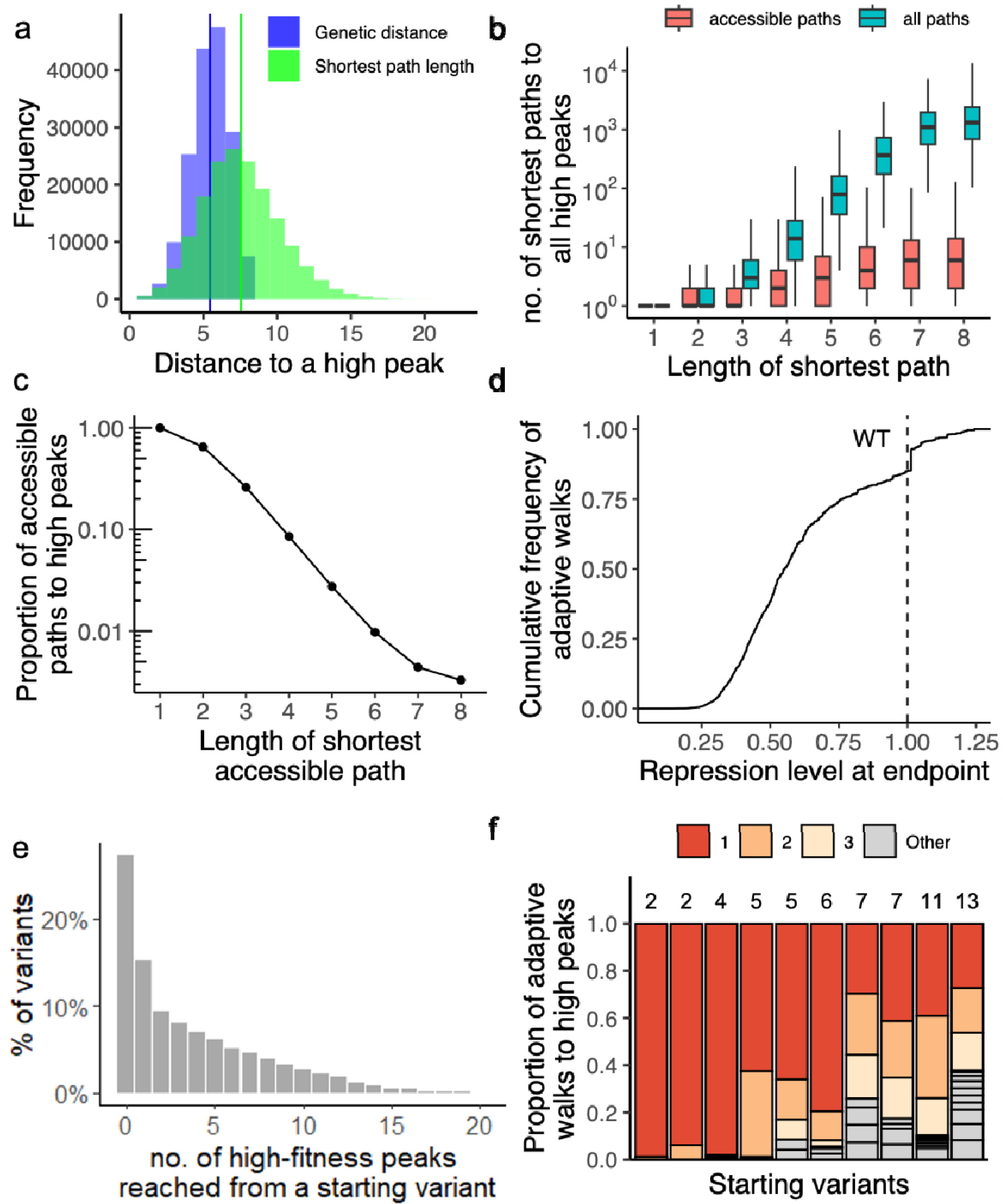
a. Accessible paths to high repression peaks tend to be short. The blue histogram shows the distribution of the genetic distances for all pairs of variants and their respective attainable peaks. The green histogram shows the distribution of the number of mutational steps for the shortest accessible paths between variants and their respective attainable peaks. **b. The landscape is abundant with paths leading to high repression peaks**. The vertical axis enumerates the count of the shortest paths for each variant towards any high peak. The horizontal axis indicates the length of the shortest paths. The red and blue boxplots represent the distribution of the shortest and accessible paths to all peaks, respectively. **c. The fraction of accessible paths decreases as path length increases**. The vertical axis displays the fraction of accessible paths (among all possible shortest paths) reaching high repression peaks. As path length increases, the proportion of accessible paths diminishes. Circles represent the average fraction of accessible paths for each path length. The vertical axis uses a logarithmic scale. **d. High repression peaks are attainable through adaptive evolution**. The panel shows the cumulative distribution of repression values reached by 10^3^ adaptive walks starting from each non-peak variant in the landscape (Methods). The dashed vertical line at *x*=1 shows the repression value of the wild type. Only 20% of adaptive walks reached a repression value of 1 or higher **e. Most starting variants can reach more than a single peak**. The bar plot shows the proportion of starting variants (vertical axis) reaching zero, one or multiple high peaks (horizontal axis). For each variant, we simulated 10^3^ populations evolving through adaptive walks. Data is based on 15,671 × 10^3^ adaptive walks, such that 10^3^ walks started from the same variant for each of 15,671 non-peak starting variants. Random walks starting from 27%, 15%, and 58% of variants reached no high peak (x=0), exactly one high peak (x=1), or more than one high peak (x>1), respectively. **f. Some high repression peaks are reached more frequently than others**. We randomly chose 10 starting variants and assessed the number of different high peaks attained by each of them. Each starting variant is represented by a bar on the horizontal axis, and the stacks represent the number of high peaks attained by that variant during its 10^3^ adaptive walks. The numbers on top of the bars represent the total number of peaks reached by each variant. Bars are ordered in ascending order along the horizontal axis according to the number of attained peaks for each variant. The height of each stack represents the proportion of adaptive walks (out of all walks attaining high peaks) attaining a specific peak, such that the sum of stack heights for a starting variant equals 1 (100%). Stacks are ordered by descending height within a bar, with the most frequently attained peak located at the top. Each variant’s three topmost accessed peaks are color-coded in red, light orange, and light yellow, respectively.

Thus far, we have defined a peak as accessible from any one variant if monotonically repression-increasing paths to it exist from the variant. However, the fraction of such accessible paths among all possible paths may be tiny, such that evolving populations may usually not find them. To find out whether this is the case, for each basin of attraction, we determined all possible paths (both accessible and inaccessible) from every single variant within that basin to its respective peak. We observed an exponential increase in the number of paths to a peak contingent upon the genetic distance of the variant to that peak (**Figure 5b**).

At short distances from a peak, a high fraction of paths is accessible, but this fraction decreases dramatically as the path length increases (**Figure 5c**). At the maximum distance of eight mutational steps, merely 1% of paths remain accessible.

### Only a minority of evolving populations reach high repression peaks

The length of accessible paths to high peaks (**Figure 5a**), their smaller numbers (**Figures 5b and 5c**), and the rarity of high repression peaks (only 3% of all peaks), raise the question of whether evolving populations would ever discover these peaks. To find out, we simulated evolution on the TetR landscape by random mutation and natural selection for strong TetR-mediated repression.

Because *E. coli* has large populations exceeding 10^8^ individuals^37^, even small effects of mutations can be detected by natural selection. In addition, the probability of mutations occurring in our 8 nucleotide positions of the wild-type binding site in the *E. coli* genome is very low (8 positions × 2.2×10^−10^ substitutions per position per generation^37^. Thus, adaptive evolution on our landscape would occur in the well-studied strong selection weak mutation regime (SSWM) ^38–41^. In this regime, any beneficial mutation is likely to become fixed before another mutant appears that will eventually go to fixation. In other words, populations are monomorphic most of the time - they occupy a single genotypic position on the landscape - and undergo an uphill walk until a peak is reached. In this regime, clonal interference between selected variants and recombination can be neglected^41,42^.

We simulated such adaptive walks as starting from each genotype in the landscape that was not a peak. We then chose at random and with uniform probability one of this genotype’s (1-mutant) neighbors with higher repression than itself as the first step in this random walk. We continued this procedure for as many steps as needed until a peak had been reached. More specifically, we performed 10^3^ simulations for each of the 15,671 non-peak starting genotypes.

Adaptive walks that terminated on high repression peaks comprised on average 6.2± 2.8 (mean±s.d.) mutational steps, 2.6 mutations more than the mean shortest genetic distance between the corresponding start and end points of the adaptive walks (mean±s.d.= 3.6± 1.5, **Supplementary Figure S14a**., Welch Two Sample t-test, t = -1259.8, df = 3603271, p-value < 2.2 × 10^−16^, N= 2,353,470). Notably, most walks reached low repression peaks, with only 20% terminating at high peaks (**Figure 5d**). This number is modest, but higher than expected when considering that only 3% of all peaks are high peaks.

Next, we wanted to find out whether population size affects this observation. In small populations, genetic drift can help populations traverse the valleys between a low peak and a higher peak, raising the question of whether small populations may attain higher peaks ^3,43^. To find out, we simulated adaptive walks with an approach pioneered by Kimura^44,45^ that allows us to calculate the fixation probability of any mutation as a function of population size (**Supplementary Methods 10**). We used this approach to simulate adaptive evolution in populations with 10^2^, 10^5^ and 10^8^ individuals, and found that 25%, 20% and 20% of walks attained high peaks, respectively (**Supplementary Figure S15**).

Finally, we also studied how deviations from the SSWM regime would affect our observations. Populations outside this regime are polymorphic most of the time^42,46,47^ and as a result, they experience clonal interference, in which the fittest of several simultaneously segregating mutations is likely to go to fixation ^42,46,47^. We simulated this scenario through “greedy” adaptive walks, in which an evolving genotype steps to the neighbor with the highest repression increase until a repression peak is reached ^5,39,48,49^. Because such walks are deterministic when started from the same genotype, we performed only one simulation for each non-peak genotype (N= 15,671). Similar to walks in the SSWM regime (**Supplementary Figure S16f**), 20% percent of populations attained a high repression peak. In sum, regardless of the regime we studied, a substantial minority of evolving populations reaches a high repression peak.

### Adaptive evolution on the TetR landscape is highly contingent on chance events

Because different peaks show overlapping basins of attraction (**Figure 4f**), adaptive evolution on the TetR landscape may be contingent on the starting genotype and on the mutational steps a population takes. Both factors may influence which peak the population reaches.

More generally, evolutionary contingency refers to the dependence of a historical process on chance events that can render this process unpredictable^50,51^. To quantify the extent of such contingency, we first simulated 1,000 SSWM adaptive walks starting from each genotype that was not a peak, assessing the number of different high peaks that can be reached from each variant.

Our findings, presented in **Figure 5d**, demonstrate that 20% of all possible walks (15,671 × 10^3^) can reach high repression peaks. However, we do not specify the number of starting variants corresponding to these walks. In **Figure 5e**, we show that of the 15,671 variants in the landscape, 73% reached at least one high peak. Combining the insights from **Figures 5d-e**, we find that the 20% of walks reaching high peaks are distributed among 73% of the starting variants.

Among starting variants from which adaptive walks attained at least one high peak, 58% reached two or more peaks (**Figure 5e**). These observations agree with our earlier observation that multiple high repression peaks are accessible from individual variants (**Figure 4d**). Not all peaks are reached with the same probability, i.e., some are reached more frequently than others (**Figure 5f**). Finally, we note that many alternative shortest paths can be assessed from a starting variant to a peak (6.37±8.43 mean±s.d, **Figure 5b**). In sum, our observations suggest that the identity of any peak attained on the TetR landscape is highly contingent on stochastic events.

## DISCUSSION

Understanding the interplay between the spaces of the “actual” – what already exists in nature – and the “possible” – what could exist – is a fundamental endeavor in evolutionary biology^52–54^. Here, we address this problem by exploring how evolution can navigate a space of possible genotypes embodied in a rationally designed library of TFBSs. We do so for the prokaryotic TF TetR^14,19^, a transcriptional repressor with only two cognate binding sequences and fewer than a dozen synthetic TFBSs ^35,55–57^.

We developed a novel plasmid-based system to characterize more TetR binding sites and study the adaptive landscape formed by these variants. Our experiments expand the available information on TetR TFBSs from less than a dozen to thousands of mutational variants.

We found that 12% of the variants in our landscape are peaks. By this measure, the TetR landscape is highly rugged. It is also rugged by another measure, a non-additive interaction between mutations known as reciprocal sign epistasis, in which two individual mutations reduce fitness (repression strength) but both mutations together increase it (**Supplementary Methods 9.5**)^58–60^. Specifically, 30% of mutant pairs in our landscape exhibit reciprocal sign epistasis (**Supplementary Figure S17, Supplementary Methods 9.5**). This is a much higher incidence than in smooth and better-studied eukaryotic regulatory landscapes, where only 4,5% of mutant interactions in TFBSs exhibit reciprocal sign epistasis^1^. Well-studied protein adaptive landscapes also show less reciprocal sign epistasis (8-22%^61–64^).

In our TFBS library, only a tiny fraction (0.44%) of peaks conveys higher repression levels than the wild type. Despite the rarity of strong repression peaks and the high ruggedness of this landscape, it also has properties typically associated with smooth landscapes. Perhaps the most striking one is that higher repression peaks are more accessible, in the sense that they have larger basins of attractions, than lower peaks. In addition, 20% or more of evolving populations reach peaks, conveying higher repression than the wild type. This holds for both large and small populations and in the presence of clonal interference (**Supplementary Methods 10**). It suggests that indicators such as the number of peaks may be misleading when studying the evolutionary accessibility of high peaks in rugged landscapes. It is also consistent with previous work showing that the relationship between peak accessibility and epistasis in complex landscapes may not be straightforward^63,65,66^.

One possible albeit speculative explanation for the navigability of the TetR binding site landscape is that it has been shaped through the evolution of TetR itself by the deleterious consequences of misregulated *tet* resistance genes. In nature, TetR regulates the expression of the TetA efflux pump^18,20^ whose loss of regulation is toxic to *E. coli*, partly because it leads to a loss of membrane potential ^19,67–69^.

Additionally, we considered all mutations in our analysis as non-neutral with respect to repression levels. When assuming extensive neutrality by simulating adaptive walks with low population sizes (N = 10^2^, **Supplementary Methods 10** and **Supplementary Figure S15**), the number of accessible paths to high peaks became even higher, because paths passing through slightly deleterious mutations became accessible. This is not surprising, because drift can help populations transverse valleys in the landscape^3,43^. In other words, our estimates of peak accessibility are conservative.

Our observation that it is easy to evolve high repression TetR binding sites raises the question of why such sites have not been observed in nature. Aside from the possibility that they remain to be discovered, there may also be a limit to which high repression may benefit TetR. For example, there may be a trade-off between the reduction of the fitness cost conferred by tight repression and the capacity of TetR to quickly respond to low concentrations of the tetracycline inducer^19,70^. Such a trade-off has been observed for the LacI repressor, which regulates the lactose-metabolizing *lac* operon. Excessive binding affinity of LacI to its TFBS requires a higher concentration of the inducer to de-repress the system^71^. If the same applies to TetR, the necessary increase of intracellular tetracycline concentration may itself be toxic. More generally, the assumption that the strongest regulation is the most beneficial may not always apply ^72–74^.

We observed extensive contingency in the adaptive evolution of high repression genotypes, because populations starting from the same low repression genotypes often reached different high repression peaks. This genotypic contingency may impact future phenotypic evolution for two reasons. First, different high repression peaks may confer varying repression strengths and, consequently, different fitness. Second, environmental variation, such as in temperature, inducer concentrations, or protein levels, may exacerbate these differences, similar to what has been observed in other bacterial transcriptional regulation systems ^70^.

Previous studies with the Lac repressor^75,76^ and the AraC ^70^ repressor demonstrated that alternating environments opened new adaptive trajectories and led to new epistatic interactions dependent on the presence of inducers. Through such environmental variation in fitness, different peaks in a landscape can become departure points for different evolutionary trajectories, further reducing the predictability of evolution^77,78^.

In sum, we introduced a new assay platform to investigate bacterial regulatory landscapes in vivo, allowing us to expand previous studies from a limited number of sequences to thousands. Our findings reveal that while the TetR landscape is highly rugged and high repression peaks are sparse, adaptive evolution can easily reach such peaks. This finding adds to the increasing body of empirical evidence suggesting a non-trivial relationship between landscape ruggedness and navigability. It also highlights a need for further refinement of current landscape theory ^63,65,66^. Additionally, our study paves the way for future studies comparing the landscapes of various transcription factors and investigating the impact of environmental changes on landscape topography and navigability.

## MATERIALS AND METHODS

Supplementary Material contains extended experimental and data analysis details.

### Strains and plasmids

We obtained electrocompetent *E. coli* cells (strain SIG10-MAX^®^) from Sigma Aldrich (CMC0004). The genotype of this strain (**Supplementary Table S2**) is similar to DH5α (Sigma Aldrich commercial information, see **Supplementary Table S2**) and is resistant to the antibiotic streptomycin. Due to its high transformation efficiency, we used it for molecular cloning, library generation, and sort-seq experiments. The design, genetic parts, and assembly of the plasmid vectors pCAW-Sort-Seq and pCAW-Sort-Seq-Neg we used in this study are available in the Supplementary material. All vectors and strains are listed in **Supplementary Tables S2 and S3**.

### Sort-seq procedure

We designed the *tetO2* variant library in the Snapgene® software (snapgene.com). It was synthesized by IDT (Coralville, USA) as a single-stranded DNA Ultramer® of 140bp (4nmol). We resuspended the library in nuclease-free distilled water and serially diluted it to a concentration of 50ng/uL. We PCR-amplified the library oligonucleotides, ligated them into the PCR-amplified pCAW-Sort-Seq plasmid backbone (low-copy number, multi-host, pBBR1 replication origin) ^79^ by Gibson assembly, and transformed cells with the resulting library after column-based purification with a commercial electroporation kit (NEB #T1020L). We grew populations of *E. coli* cells hosting the library in LB medium supplemented with 50 μg/mL chloramphenicol to saturation (overnight growth, 16 hours, 200 rpm, 37°C). After overnight growth, we diluted the overnight cultures in LB medium supplemented with 50 μg/mL chloramphenicol in a 1:100 ratio (v/v) and grew the cultures for 5 h until they reached late-exponential/early -stationary phase. Before sorting, we diluted 50μL of the cultures in 1mL of cold Dulbecco’s PBS (Sigma-Aldrich #D8537) in 15 mL FACS tubes. We performed FACS-sorting on a FACS Aria III flow cytometer (BD Biosciences, San Jose, CA). We sorted and binned cells according to their GFP-mediated fluorescence (FITC-H channel, 488nm laser, emission filters 502LP, 530/30). First, we recorded the fluorescence of 10^6^ cells and created 13 evenly spaced gates (“bins”) on a binary logarithmic (log_2_) scale spanning the range of fluorescence values. Because logarithmically (log_2_) spaced fluorescence intervals provide a reliable resolution for sort-seq studies^23,80^, we subdivided the range of observed fluorescence values in our library into 13 such intervals or “bins” (**see methods, Supplementary Methods 7, and Supplementary Figures S4-S5**). The number of cells sorted into each bin corresponded to the fraction of the 10^6^ cells recorded for that bin. Thus, the total number of sorted cells per replicate was 10^6^. We isolated plasmids from cells sorted into each bin (Qiagen, Germany) after overnight growth, and used a polymerase chain reaction (PCR) to amplify the *tetO2* region from each plasmid for Illumina sequencing. The distribution of variants among the different bins allowed us to map each individual genotype to a repression level (see **Supplementary Methods: *9*.*2*, Supplementary Figures S9-S10**). The expression values of cells in each bin were reproducible, as confirmed by regrowth and re-measuring fluorescence distributions (**Supplementary Methods 7, and Supplementary Figure S5**). The Supplementary material has additional details on library construction and sort-seq procedures.

### Replicates

Data from high-throughput methods to measure gene expression are intrinsically noisy. The reasons include biological cell-to-cell expression variation ^81^, as well as technical ^82–85^ variation (e.g. pipetting errors, equipment biases, limit of detection thresholds etc.). In order to account for such variability, we carried out the sorting procedure in three replicates derived from three independent library transformations.

### Library diversity

The resulting library contained 17,851 genotypes, representing 27% of the 4^8^ genotypes in the genotype space we study. We did not recover all genotypes because of two common technical constraints in mutational library studies: biased PCR amplification ^86^ and loss of sequence diversity after sorting cells ^25^. Moreover, the inclusion of individual sequence variants for analysis met a stringent sequencing depth threshold to reduce false-positive reads (**Supplementary Methods 7, Supplementary Figure S8**).

### Repression levels

Due to gene expression and measurement noise, individual TFBS variants in a sort-seq experiment usually appear in more than a single bin, and their read count (frequency) varies among bins^23,87–89^ (**Supplementary Figures S9-S10**). Following established practice^80,88^, we used a weighted average of these frequencies for each variant to represent the mean expression level driven by the variant (**Supplementary Methods 9.2**). To facilitate interpretation, we converted this expression level into a repression level and normalized it to the repression of the wild type (**Supplementary Methods 9.2**). On this scale, the repression level of the wild type has a value of 1. Values below one indicate weaker repression (weaker TetR binding, higher GFP expression), whereas values above one indicate stronger repression (lower GFP expression).

### Validating repression levels with plate reader measurements

To further validate our calculated repression levels, we plated cells from each fluorescence bin onto LB-agar plates, picked 15 colonies of cells from each bin (N=195), sequenced their TFBS sequence and replated them onto LB-agar plates. We picked individual colonies and grew them overnight (16 hours, 37°C, 220 rpm) in liquid LB supplemented with 50 μg/mL of chloramphenicol. We diluted the cultures to 1:10 (v/v) in cold Dulbecco’s PBS (Sigma-Aldrich #D8537) to a final volume of 1 mL. We transferred 200 μl of the diluted cultures into individual wells in 96-well plates and measured GFP fluorescence (emission: 485nm/excitation: 510nm, bandpass: 20nm, gain: 50) as well as OD_600_. We then normalized fluorescence by OD_600_ measurements, and compared the obtained ratios to the previously calculated repression levels for the 195 selected variants. We observed a strong negative Pearson correlation (*R*=-0.86, **Figure 2c**). We performed all such measurements in biological and technical triplicates (three colonies per sample and three wells per colony, respectively).

### Code Availability and Data Analysis

All code used for processing data and plotting as well as the final processed data, plasmid sequences, and primer sequences are available on our GitHub repository (https://github.com/Cauawest; DOI: **X**). Sort-Seq sequencing files are available at the Sequence Read Archive (accession no.**X**).

**Supplementary Information** is linked to the online version of the paper

## Acknowledgements

We would like to acknowledge financial support by Swiss National Science Foundation grant 310030_208174. We would also like to thank the UZH University Priority Research Program in Evolutionary Biology, the UZH flow cytometry facility, and the Functional Genomics Center Zurich for technical support. We thank Diego Pesce and Andrei Papkou for help in establishing experimental protocols, computational analysis, as well as for theoretical discussions.

## Author contributions

C.A.W. and A.W conceived the study and designed experiments.

C.A.W. carried out experiments. C.A.W and A.W. analyzed data.

C.A.W wrote computer code to carry out bioinformatic work and analysis. C.A.W generated figures. L.G. wrote computer code to carry out simulations. C.A.W. and A.W. wrote the paper, which was edited by all authors.

## Data availability

The data generated in this study will be deposited in a public repository upon manuscript acceptance.

## Computer code

The computer code will be shared in a public repository upon manuscript acceptance.

## Competing interests

The authors declare no competing interest

## Notes

### Competing Interest Statement

The authors have declared no competing interest.

## REFERENCES

1. Aguilar-Rodríguez, J., Payne, J. L. & Wagner, A. A thousand empirical adaptive landscapes and their navigability. Nat Ecol Evol 1, 0045 (2017).

2. Fragata, I., Blanckaert, A., Dias Louro, M. A., Liberles, D. A. & Bank, C. Evolution in the light of fitness landscape theory. Trends Ecol Evol 34, 69–82 (2019).

3. Wright, S. The roles of mutation, inbreeding, crossbreeding and selection in evolution. Proc of the 6th International Congress of Genetics Preprint at (1932).

4. Kauffman, S. & Levin, S. Towards a general theory of adaptive walks on rugged landscapes. J Theor Biol 128, 11–45 (1987).

5. Kauffman, S. & Levin, S. Towards a general theory of adaptive walks on rugged landscapes. J Theor Biol 128, 11–45 (1987).

6. Wray, G. A. The evolutionary significance of cis-regulatory mutations. Nat Rev Genet 8, 206–216 (2007).

7. Hill, M. S., vande Zande, P. & Wittkopp, P. J. Molecular and evolutionary processes generating variation in gene expression. Nature Reviews Genetics vol. 22 203–215 Preprint at 10.1038/s41576-020-00304-w (2021).

8. Wittkopp, P. J. & Kalay, G. Cis-regulatory elements: Molecular mechanisms and evolutionary processes underlying divergence. Nat Rev Genet 13, 59–69 (2012).

9. Coulon, A., Chow, C. C., Singer, R. H. & Larson, D. R. Eukaryotic transcriptional dynamics: from single molecules to cell populations. Nature Reviews Genetics 2013 14:8 14, 572–584 (2013).

10. Browning, D. F. & Busby, S. J. W. Local and global regulation of transcription initiation in bacteria. Nat Rev Microbiol 14, 638–650 (2016).

11. Payne, J. L. & Wagner, A. The Robustness and Evolvability of Transcription Factor Binding Sites. Science (1979) 343, (2014).

12. Schweizer, G. & Wagner, A. Both Binding Strength and Evolutionary Accessibility Affect the Population Frequency of Transcription Factor Binding Sequences in Arabidopsis thaliana. Genome Biol Evol 13, (2021).

13. Aguilar-Rodríguez, J. & Payne, J. L. Robustness and Evolvability in Transcriptional Regulation. Evolutionary Systems Biology 197–219 (2021) doi:10.1007/978-3-030-71737-7_9.

14. Ramos, J. L. et al. The TetR family of transcriptional repressors. Microbiol Mol Biol Rev 69, 326–56 (2005).

15. Orth, P., Schnappinger, D., Hillen, W., Saenger, W. & Hinrichs, W. Structural basis of gene regulation by the tetracycline inducible Tet repressor-operator system. Nat Struct Biol 7, 215–219 (2000).

16. Herrin, G. L., Russell, D. R. & Bennett, G. N. A stable derivative of pBR322 conferring increased tetracycline resistance and increased sensitivity to fusaric acid. Plasmid 7, 290–293 (1982).

17. Bertram, R. & Hillen, W. The application of Tet repressor in prokaryotic gene regulation and expression. Microb Biotechnol 1, 2 (2008).

18. Meier, I., Wray, L. V. & Hillen, W. Differential regulation of the Tn10-encoded tetracycline resistance genes tetA and tetR by the tandem tet operators O1 and O2. EMBO J 7, 567–572 (1988).

19. Berens, C. & Hillen, W. Gene regulation by tetracyclines. Constraints of resistance regulation in bacteria shape TetR for application in eukaryotes. Eur J Biochem 270, 3109–3121 (2003).

20. Hillen, W. & Berens, C. MECHANISMS UNDERLYING EXPRESSION OF TN10 ENCODED TETRACYCLINE RESISTANCE. 10.1146/annurev.mi.48.100194.002021 48, 345–369 (2003).

21. Lutz, R. & Bujard, H. Independent and tight regulation of transcriptional units in Escherichia coli via the LacR/O, the TetR/O and AraC/I1-I2 regulatory elements. Nucleic Acids Res 25, 1203–10 (1997).

22. Kleinschmidt, C., Tovar, K., Hillen, W. & Porschke, D. Dynamics of Repressor-Operator Recognition: The Tn 70-Encoded Tetracycline Resistance Control1”. Nucleic Acids Res 27, 105–118 (1094).

23. Peterman, N. & Levine, E. Sort-seq under the hood: Implications of design choices on large-scale characterization of sequence-function relations. BMC Genomics 17, 1–17 (2016).

24. Kinney, J. B. & McCandlish, D. M. Massively Parallel Assays and Quantitative Sequence–Function Relationships. Annu Rev Genomics Hum Genet 20, annurevgenom-083118-014845 (2019).

25. Kinney, J. B., Murugan, A., Callan, C. G. & Cox, E. C. Using deep sequencing to characterize the biophysical mechanism of a transcriptional regulatory sequence. Proceedings of the National Academy of Sciences 107, 9158–9163 (2010).

26. Gerland, U. & Hwa, T. On the Selection and Evolution of Regulatory DNA Motifs. J Mol Evol 55, 386–400 (2002).

27. Mustonen, V., Kinney, J., Callan, C. G. & Lassig, M. Energy-dependent fitness: A quantitative model for the evolution of yeast transcription factor binding sites. Proceedings of the National Academy of Sciences (2008) doi:10.1073/pnas.0805909105.

28. Lynch, M. & Hagner, K. Evolutionary meandering of intermolecular interactions along the drift barrier. Proceedings of the National Academy of Sciences (2014) doi:10.1073/pnas.1421641112.

29. Mustonen, V. & Lassig, M. Evolutionary population genetics of promoters: Predicting binding sites and functional phylogenies. Proceedings of the National Academy of Sciences 102, 15936–15941 (2005).

30. Shultzaberger, R. K., Malashock, D. S., Kirsch, J. F. & Eisen, M. B. The Fitness Landscapes of cis-Acting Binding Sites in Different Promoter and Environmental Contexts. PLoS Genet 6, e1001042 (2010).

31. van Hijum, S. A. F. T., Medema, M. H. & Kuipers, O. P. Mechanisms and Evolution of Control Logic in Prokaryotic Transcriptional Regulation. Microbiology and Molecular Biology Reviews 73, 481–509 (2009).

32. Balleza, E. et al. Regulation by transcription factors in bacteria: Beyond description. FEMS Microbiol Rev 33, 133–151 (2009).

33. Price, M. N., Wetmore, K. M., Deutschbauer, A. M. & Arkin, A. P. A Comparison of the Costs and Benefits of Bacterial Gene Expression. PLoS One 11, e0164314 (2016).

34. Bank, C. Epistasis and Adaptation on Fitness Landscapes. 10.1146/annurev-ecolsys-102320-112153 53, 457–479 (2022).

35. Bolintineanu, D. S. et al. Investigation of changes in tetracycline repressor binding upon mutations in the tetracycline operator. J Chem Eng Data 59, 3167–3176 (2014).

36. Szendro, I. G., Schenk, M. F., Franke, J., Krug, J. & De Visser, J. A. G. M. Quantitative analyses of empirical fitness landscapes. Journal of Statistical Mechanics: Theory and Experiment 2013, P01005 (2013).

37. Lynch, M. et al. Genetic drift, selection and the evolution of the mutation rate. Nature Reviews Genetics 2016 17:11 17, 704–714 (2016).

38. Bank, C., Matuszewski, S., Hietpas, R. T. & Jensen, J. D. On the (un)predictability of a large intragenic fitness landscape. Proc Natl Acad Sci U S A 113, 14085–14090 (2016).

39. Orr, H. A. The population genetics of adaptation: the adaptation of DNA sequences. Evolution 56, 1317–1330 (2002).

40. Vaishnav, E. D. et al. The evolution, evolvability and engineering of gene regulatory DNA. Nature 2022 603:7901 603, 455–463 (2022).

41. Gillespie, J. H. MOLECULAR EVOLUTION OVER THE MUTATIONAL LANDSCAPE. Evolution (N Y) 38, 1116–1129 (1984).

42. Li, J., Amado, A. & Bank, C. Rapid adaptation of recombining populations on tunable fitness landscapes. Mol Ecol (2023) doi:10.1111/MEC.16900.

43. Weissman, D. B., Desai, M. M., Fisher, D. S. & Feldman, M. W. The rate at which asexual populations cross fitness valleys. Theor Popul Biol 75, 286–300 (2009).

44. Kimura, M. ON THE PROBABILITY OF FIXATION OF MUTANT GENES IN A POPULATION. Genetics 47, 713–719 (1962).

45. Crow, J. and Kimura, M. An Introduction to Population Genetics Theory [Paperback]. 608 (2009).

46. Melissa, M. J., Good, B. H., Fisher, D. S. & Desai, M. M. Population genetics of polymorphism and divergence in rapidly evolving populations. Genetics 221, (2022).

47. Stolyarova, A. V. et al. Complex fitness landscape shapes variation in a hyperpolymorphic species. Elife 11, (2022).

48. Park, S.-C., Neidhart, J. & Krug, J. Greedy adaptive walks on a correlated fitness landscape. J Theor Biol 397, 89–102 (2016).

49. Orr, H. A. A Minimum on the Mean Number of Steps Taken in Adaptive Walks. J Theor Biol 220, 241–247 (2003).

50. Gould, S. J. Wonderful Life; The Burgess Shale and the Nature of History. J Hist Biol 24, 163–170 (1992).

51. Blount, Z. D., Lenski, R. E. & Losos, J. B. Contingency and determinism in evolution: Replaying life’s tape. Science (1979) 362, (2018).

52. Hochberg, M. E., Marquet, P. A., Boyd, R. & Wagner, A. Innovation: an emerging focus from cells to societies. Philosophical Transactions of the Royal Society B: Biological Sciences 372, 20160414 (2017).

53. François Jacob. The Possible and the Actual. (Pantheon, 1982).

54. Dennett, D. C. Darwin’s Dangerous Idea: Evolution and the Meanings of Life. (Simon & Schuster, 1995).

55. Krueger, M., Scholz, O., Wisshak, S. & Hillen, W. Engineered Tet repressors with recognition specificity for the tetO-4C5G operator variant. Gene 404, 93–100 (2007).

56. Helbl, V. & Hillen, W. Stepwise selection of TetR variants recognizing tet operator 4C with high affinity and specificity. J Mol Biol 276, 313–318 (1998).

57. Helbl, V., Tiebel, B. & Hillen, W. Stepwise selection of TetR variants recognizing tet operator 6C with high affinity and specificity. J Mol Biol 276, 319–324 (1998).

58. Poelwijk, F. J., Tnase-Nicola, S., Kiviet, D. J. & Tans, S. J. Reciprocal sign epistasis is a necessary condition for multi-peaked fitness landscapes. J Theor Biol 272, 141–144 (2011).

59. Saona, R., Kondrashov, F. A. & Khudiakova, K. A. Relation Between the Number of Peaks and the Number of Reciprocal Sign Epistatic Interactions. Bull Math Biol 84, (2022).

60. Kvitek, D. J. & Sherlock, G. Reciprocal sign epistasis between frequently experimentally evolved adaptive mutations causes a rugged fitness landscape. PLoS Genet (2011) doi:10.1371/journal.pgen.1002056.

61. Li, C., Qian, W., Maclean, C. J. & Zhang, J. The fitness landscape of a tRNA gene. Science (1979) (2016) doi:10.1126/science.aae0568.

62. Li, C. & Zhang, J. Multi-environment fitness landscapes of a tRNA gene. Nature Ecology & Evolution 2018 2:6 2, 1025–1032 (2018).

63. Papkou, A., Garcia-Pastor, L., Escudero, J. A. & Wagner, A. A rugged yet easily navigable fitness landscape of antibiotic resistance. bioRxiv 2023.02.27.530293 (2023) doi:10.1101/2023.02.27.530293.

64. Song, S. & Zhang, J. Unbiased inference of the fitness landscape ruggedness from imprecise fitness estimates. Evolution (N Y) 75, 2658–2671 (2021).

65. Lagator, M., Sarikas, S., Acar, H., Bollback, J. P. & Guet, C. C. Regulatory network structure determines patterns of intermolecular epistasis. Elife 6, 1–22 (2017).

66. Greenbury, S. F., Louis, A. A. & Ahnert, S. E. The structure of genotype-phenotype maps makes fitness landscapes navigable. Nature Ecology & Evolution 2022 6:11 6, 1742–1752 (2022).

67. Eckert, B. & Beck, C. F. Overproduction of transposon Tn10-encoded tetracycline resistance protein results in cell death and loss of membrane potential. J Bacteriol 171, 3557–3559 (1989).

68. Nguyen, T. N. M., Phan, Q. G., Duong, L. P., Bertrand, K. P. & Lenski, R. E. Effects of carriage and expression of the Tn10 tetracycline-resistance operon on the fitness of Escherichia coli K12. Mol Biol Evol 6, 213–225 (1989).

69. Rajer, F. & Sandegren, L. The Role of Antibiotic Resistance Genes in the Fitness Cost of Multiresistance Plasmids. mBio 13, (2022).

70. Lagator, M., Igler, C., Moreno, A. B., Guet, C. C. & Bollback, J. P. Epistatic interactions in the arabinose cis-regulatory element. Mol Biol Evol 33, 761–769 (2016).

71. Razo-Mejia, M. et al. Tuning Transcriptional Regulation through Signaling: A Predictive Theory of Allosteric Induction. Cell Syst 6, 456–469.e10 (2018).

72. Srivastava, M. & Payne, J. L. On the incongruence of genotype-phenotype and fitness landscapes. PLoS Comput Biol 18, e1010524 (2022).

73. de Visser, J. A. G. M., Cooper, T. F. & Elena, S. F. The causes of epistasis. Proceedings of the Royal Society B: Biological Sciences 278, 3617–3624 (2011).

74. Crocker, J., Preger-Ben Noon, E. & Stern, D. L. The Soft Touch: Low-Affinity Transcription Factor Binding Sites in Development and Evolution.Current Topics in Developmental Biology vol. 117 (Elsevier Inc., 2016).

75. Poelwijk, F. J., Kiviet, D. J., Weinreich, D. M. & Tans, S. J. Empirical fitness landscapes reveal accessible evolutionary paths. Nature (2007) doi:10.1038/nature05451.

76. de Vos, M. G. J., Poelwijk, F. J., Battich, N., Ndika, J. D. T. & Tans, S. J. Environmental Dependence of Genetic Constraint. PLoS Genet 9, e1003580 (2013).

77. Majic, P. The Molecular Scaffolds of the élan vital. Parrhesia: A Journal of Critical Philosophy (2022).

78. Lässig, M., Mustonen, V. & Walczak, A. M. Predicting evolution. Nature Ecology & Evolution 2017 1:3 1, 1–9 (2017).

79. Jahn, M., Vorpahl, C., Hübschmann, T., Harms, H. & Müller, S. Copy number variability of expression plasmids determined by cell sorting and droplet digital PCR. Microb Cell Fact 15, 211 (2016).

80. Lagator, M. et al. Predicting bacterial promoter function and evolution from random sequences. Elife 11, (2022).

81. Elowitz, M. B., Levine, A. J., Siggia, E. D. & Swain, P. S. Stochastic Gene Expression in a Single Cell. Science (1979) 297, 1183–6 (2007).

82. Beal, J. et al. Quantification of bacterial fluorescence using independent calibrants. PLoS One 13, e0199432 (2018).

83. Beal, J. et al. Reproducibility of fluorescent expression from engineered biological constructs in E. coli. PLoS One 11, (2016).

84. Beal, J., Haddock-Angelli, T., Farny, N. & Rettberg, R. Time to Get Serious about Measurement in Synthetic Biology. Trends in Biotechnology (2018) doi:10.1016/j.tibtech.2018.05.003.

85. Trippe, B. L. et al. Randomized gates eliminate bias in sort-seq assays. Protein Science 31, e4401 (2022).

86. Kebschull, J. M. & Zador, A. M. Sources of PCR-induced distortions in high-throughput sequencing data sets. Nucleic Acids Res 43, e143–e143 (2015).

87. Gilliot, P.-A. & Gorochowski, T. E. Effective design and inference for cell sorting and sequencing based massively parallel reporter assays. bioRxiv 2022.11.07.515414 (2022) doi:10.1101/2022.11.07.515414.

88. de Boer, C. G. et al. Deciphering eukaryotic gene-regulatory logic with 100 million random promoters. Nature Biotechnology 2019 38:1 38, 56–65 (2019).

89. Belliveau, N. M. et al. A Systematic and Scalable Approach for Dissecting the Molecular Mechanisms of Transcriptional Regulation in Bacteria. Biophys J 114, 151a (2018).

