## Supplementary material for "The highly rugged yet navigable regulatory landscape of the bacterial transcription factor TetR": Suppl_Material

**SUPPLEMENTARY METHODS, TABLES AND FIGURES**

**Index**

**Supplementary Methods**

1. **Media and reagents**
2. **General cloning procedures**
   2.1. *Overnight incubation of cultures in liquid and solid medium*
   2.2. *PCR Reactions*
   2.3. *Verifying PCR products through gel electrophoresis*
   2.4. *DNA purification with commercial kits*
   2.5. *DNA purification through ethanol precipitation*
   2.6. *Gibson assembly*
   2.7. *Preparation of electro-competent cells*
   2.8. *Electroporation*
3. **The pCAW-Sort-Seq design**
4. **Assembling the pCAW-Sort-Seq plasmid**
5. **Library design, synthesis and amplification:**
6. **Library cloning**
7. **Analysing and sorting cells**
8. **DNA extraction and DNA preparation for sequencing**
9. **Data analysis**
   9.1. *Filtering and preparing sequencing reads*
   9.2. *Calculating repression levels*
   9.3. *Combining the data from triplicates*
   9.4. *Frequency matrix and sequence logo*
   9.5. *Creation of genotype networks and network metrics*
   9.6. *Principal component analysis*
10. **Simulated adaptive walks**

**Supplementary Tables**

**S1. Network metrics**

**S2. Table of strains**

**S3. Table of plasmids**

**S4. Primers for engineering the pCAW-sort-seq plasmid**

**S5. Primers for constructing libraries**

**S6. Primers with barcodes for sequencing bins**

**Supplementary Figures**

**S1. The pCAW-SortSeq-TetR plasmid.**

**S2. The repression measurement module of the pCAW-SortSeq-TetR plasmid**

**S3 Validating the pCAW-SortSeq-TetR plasmid with *tetO2* mutants**

**S4. Flow cytometry gating and fluorescence expression levels for controls and library**

**S5. Distribution of fluorescence levels after sorting of cells with the library into bins.**

**S6. Distribution of sequence reads per bins**

**S7. Correlation of read coverage in the *tetO2* mutant library**

**S8. Number of genotypes obtained by choosing different sequencing depth (read count) cutoffs**

**S9. Genotypes per fluorescence bin and overlaps between them**

**S10. Number of bins per genotype**

**S11. DNA Sequence LOGO obtained by a previous study**

**S12. Nucleotide contribution in principal component analysis**

**S13. The distribution of basin sizes among peaks**

**S14. Adaptive walks in which each mutational step is chosen with uniform probability among all repression-increasing steps**

**S15. Adaptive walks using the Kimura model with different population sizes**

**S16. Greedy adaptive walk simulations**

**S17. Epistatic interactions**

**Supplementary Methods**

**1	Media and reagents**

To prepare SOB medium we dissolved 25.5g of its solid stock (VWR J906) in 960 ml of distilled water and autoclaved the medium before use. To prepare SOC medium, we added 20 ml of 1 M D-glucose (Sigma G8270), and 20 ml of 1 M magnesium sulfate (Sigma 230391) to 960 ml of SOB medium. To prepare LB medium we dissolved 25g of its solid stock (Sigma-Aldrich L3522) in 1 liter of distilled water and autoclaved the medium before use. We purchased M9 minimal salt from Sigma (M6030), dissolved it according to the supplier’s instructions, sterilized the solution by autoclaving, and supplemented it with 0.4% glucose (Sigma G8270), 0.2% casamino acid (Merk Millipore, 2240), 2 mM magnesium sulfate (Sigma 230391), and 0.1 mM calcium chloride (Sigma C7902).Where necessary, we supplemented growth media with chloramphenicol (50 µg/mL working concentration) and/or anhydrotetracycline (100 ng/mL working concentration) (Cayman-chemicals #10009542).

**2.	General procedures:**

*2.1	Overnight incubation of cultures in liquid and solid medium*

Unless otherwise stated, we grew overnight liquid cultures on liquid LB medium (15mL or 50mL Falcon tubes) supplemented with chloramphenicol (50 µg/mL working concentration) for 16 hours at 37°C and 200rpm (50 mm orbital motion) on an Infors HT Multitron Incubator Shaker. Unless otherwise stated, we incubated bacterial colonies on solid LB-agar medium (sterile 90mm × 15mm plastic petri dishes) supplemented with chloramphenicol (50 µg/mL working concentration) for 16 hours at 37°C.

*2.2	PCR Reactions*

Unless otherwise stated, we amplified DNA fragments by PCR using a Q5^®^ high-fidelity polymerase (NEB #M0491L) to reduce the probability of introducing mutations into the amplicons. We adopted the reaction protocol provided by NEB for a final volume of 50uL. We performed each reaction in duplicate and pooled the reaction products at the end of the PCR program. We calculated the primer melting temperatures (Tm) following the NEB Tm calculator () for a primer concentration of 500nM.

*2.3	Verifying PCR products through gel electrophoresis*

Unless otherwise stated, after PCR amplification, we checked reactions for the presence of single-band amplicons (and the absence of unspecific bands) through electrophoresis. We performed electrophoresis in a 0.8% agarose Tris-EDTA (TAE) gel for 45 minutes at 120V, or until the gel bands had migrated more than halfway across the total length of the gel.

*2.4	DNA purification with commercial kits*

After confirming the presence of single bands during electrophoresis, we purified the samples using the Monarch® DNA PCR/Gel Extraction Kit (NEB #T1020L), following the original protocol provided with the kit. When needed (e.g., in the presence of unspecific bands after PCR), we gel purified samples with the following protocol. We added 10 μL of 6x NEB DNA dye to each 50 μL PCR reaction, loaded the full volumes on a 1% agarose gel, and performed electrophoresis for 45 minutes at 120V, or until the gel bands had migrated more than halfway across the total length of the gel. We used a scalpel to remove the DNA band corresponding to the amplified sequence. We performed the gel extraction using the Monarch® DNA Gel Extraction Kit (NEB #T1020L).

*2.5	DNA purification through ethanol precipitation*

We precipitated DNA by adding 1 µL of glycogen (R0551, Thermo Scientific) to 20 µL of to the inactivated ligation reaction, 50 µL of 7.5 M ammonium acetate (A2706-100ML, Sigma), 375 µL of ice-cold absolute ethanol, and 80 µL of ddH2O. After incubating at -20°C for 20 min, we centrifuged the mixture at 18,000 g for 20 min at 4°C. We washed the precipitate twice using 800 µL of cold ethanol (70%). After drying the precipitate using an Eppendorf concentrator 5301, we dissolved it in at least 10 µL of ddH2O.

*2.6	Gibson assembly:*

We performed the Gibson assembly of PCR amplified fragments with an NEBuilder-HiFi® DNA Assembly Master Mix kit (NEB #E2621L). We calculated molarity for the assembly based on the protocol provided by the Barrick Lab (<https://barricklab.org/twiki/bin/view/Lab/ProtocolsGibsonCloning>). We incubated the reaction mixture for 1 hour at 50°C in a dry bath incubator, and then placed it on ice for further use.

*2.7	Preparation of electro-competent cells*

We prepared electro-competent cells using glycerol/mannitol step centrifugation^1^ . Briefly, we grew the chosen *E. coli* strains in 5 mL SOB medium at 37°C and 250 rpm overnight. We transferred 3 mL culture into 300 mL SOB medium the next morning and continued to incubate the transferred culture at 37°C and 250 rpm until its OD600 had reached a value between 0.4 and 0.6 (optical path length: 1 cm, 2-4 hours). We cooled the culture on ice for 15 min and collected cells at 4°C by centrifuging at 1,500 g for 15 min. We used 60 mL ice-cold ddH2O to suspend the cells and distributed them into three 50 mL tubes. Then we slowly added 10 mL ice cold glycerol/mannitol solution (20% glycerol (w/v) and 1.5% mannitol (w/v)) to the bottom of each tube by using a 10 mL pipette. We centrifuged the tubes at 1,500 g and 4°C for 15 min in a centrifuge (Eppendorf 5810/5810 R) by setting acceleration/deceleration to zero. We removed the supernatant and suspended the cells in 3.0 mL ice-cold glycerol/mannitol solution. Subsequently, we transferred 100 µL of the resulting suspensions into pre-cooled 1.5 mL tubes and incubated them in a dry ice-ethanol bath for ~1 min. Then we stored the suspensions at -80°C for transformation experiments.

*2.8	Electroporation*

For all the transformation procedures in this work, we transformed 100 µL of electrocompetent cells by electroporation in 0.2 cm cuvettes (EP202, Cell Projects, UK), through a Micropulser electroporator (Bio-Rad) set at EC3 (15k V/cm). We recovered electroporated cells in 1mL of pre-warmed SOC media in 15mL falcon tubes for 1,5 hours (37°C, 220 rpm). Except when stated otherwise, we plated 300 µL of the recovered culture on an LB agar plate supplemented with 50 μg/mL of chloramphenicol and incubated overnight (16 hours) at 37 °C for subsequent screening and confirmation of clones through Sanger sequencing.

**3.	The pCAW-Sort-Seq design**

We designed the pCAW-Sort-Seq plasmid in the Snapgene® software ([www.snapgene.com](https://www.snapgene.com/)) by combining information from different literature sources. Below, we provide a more detailed explanation of the plasmid’s components.

The pCAW-Sort-Seq plasmid (**Figure 1b**, **Supplementary Figures S1-S2**) is a PSEVA231-derived plasmid ^2^, harbouring a pBBR1 replication origin (broad-host, low copy, i.e., 5 to 10 copies per cell) ^3^ and a chloramphenicol resistance gene. We chose the pBBR1 origin of replication for two main reasons. First, a low-copy system is better suited for simulating the native protein-DNA interactions observed in the single-copy bacterial chromosome. Second, our system is compatible with a wide range of gram-negative bacteria, allowing the generation of genotype-phenotype maps in different bacterial hosts. The TetR repression system we adopted in this work is based on ref.^4^. We use anhydrotetracycline (Cayman-chemicals#10009542) as an inducer to derepress the system when required. We diluted anhydrotetracycline in absolute ethanol in a stock concentration (1000X) of 100 µg/mL to a working concentration of 100 ng/mL.

The fluorescent reporter gene of the pCAW-Sort-Seq plasmid is *sfgfp*^5^. It provides information on TetR repression/binding strength. The *tetr* gene is constitutively expressed under a low-strength pLac promoter variant developed by ^6^. Measuring TetR repression through sfGFP fluorescence relies on having a *tetO2* (5’-TCCCTATCAGTGATAGAGA-3’) binding site between a medium strength promoter (the BBa_J23110 promoter from <http://parts.igem.org/Part:BBa_J23110> ^7^) and the *sfgfp* gene, at the +10 position relative to the transcription starting site (TSS) of the gene ^8^. The region where the *tetO2* binding site is located can be easily replaced for generating TFBS libraries through restriction digestion (HindIII and BamHI restriction sites) or Gibson assembly^9^. The reporter gene *sfgfp* is insulated by a transcriptional insulator named RiboJ, a synthetic ribozyme that removes 5’UTR interferences from variable TFBS sequence in the mRNA by self-cleavage^10^. Promoters are preceded by insulator sequences, terminators that alleviate contextual effects of upstream sequences over promoter regions^11^. We based the divergent orientation of regulatory modules on ref. ^12^. We retrieved natural (*ECK*) and synthetic transcriptional terminators (*synterm*) from ref. ^13^. Our system is also compatible with barcoding for *in vivo* multiplexed measurements based on RNA-Seq, as proposed by ^14^. We analyzed the plasmid with the EFM calculator ^15^ for genetic stability, and iteratively modified its design until a low instability score (RIP Score: 50.0) was reached.

**4.	Assembling the pCAW-Sort-Seq plasmid**

The designed plasmid was synthesized by Twist Biosciences (USA) in two fragments that we joined through Gibson assembly ^16^ using the pCAW_frag1_F/pCAW_frag1_R, pCAW_frag2_F/pCAW_frag2_R primer sets (**Supplementary Table S4**). For a more in-depth view of the plasmid map, see **Supplementary Figure S1**, or the Addgene collection browser (<https://www.addgene.org/>), ID (to be uploaded).

We amplified the synthesized DNA fragments by PCR, performing each reaction in duplicate, and pooled the reaction products at the end of the PCR. We performed PCR with the following program: 98 °C/30 s; 25 cycles of 98 °C/10 s, 64 °C/30 s, 72 °C/30 s; 72 °C/2 min.

After the PCR amplification, we checked reactions for the presence of single-band amplicons and the absence of unspecific bands through gel electrophoresis. After confirming the presence of single bands, we treated the pooled PCR reactions with 2 µL of DpnI restriction enzyme (NEB #R0176L), and incubated them at 37 °C for 2 hours to remove traces of the template plasmid. We purified digested samples using the Monarch® DNA gel extraction kit (NEB #T1020L). We then performed the assembly of the two fragments with a NEBuilder-HiFi® DNA Assembly Master Mix kit (NEB #E2621L). We calculated molarity for the assembly based on the protocol provided by the Barrick Lab (<https://barricklab.org/twiki/bin/view/Lab/ProtocolsGibsonCloning>).

Subsequently, we combined an equimolar ratio of the two fragments to a total volume of 5µL, and added 5 µL of the Gibson master mix (2X). We incubated the resulting 10 µL reaction at 50°C for 1 hour in a dry-bath incubator. For transformation, we directly added 1 µL of the reaction to 100 µL of electrocompetent cells, and performed transformation as described in the section *Electroporation*. When needed, we purified the reaction using the Monarch® DNA gel extraction kit (NEB #T1020L) prior to transformation, resuspended the purified DNA in 10 µL of nuclease-free distilled water, and used all 10 µL for transformation. After transformation, we plated recovered cultures on LB agar plates and incubated them at 37°C overnight. We picked colonies and directly sent them for clone confirmation through Sanger sequencing (NightSeq® service, Microsynth, Switzerland). We grew individual clones overnight in LB liquid media supplemented with chloramphenicol, aliquoted them into 1mL cryotubes as glycerol stocks (glycerol (20% v/v final concentration) and stored them at -80°C for further use.

**5.	Library design, synthesis and amplification:**

Several studies have identified the most important positions for *tetO2* DNA binding ^4,17^. Based on these studies, we designed the *tetO2*-derived library by randomizing the eight most important positions. The randomized sites form two symmetric palindromes of four base pairs each. The calculated library size for this approach is (4^8^)^=^65,536 sequences. The originally published *tetO2*-original sequence ^4^ is 5’-TCCCTATCAGTGATAGAGA-3’. Our *tetO2*-derived library sequence is 5’-TCCCNNNNAGTNNNNGAGA-3’, where N represents the randomized positions.

We designed the library in the Snapgene® software (snapgene.com), and had it synthesized by IDT (Coralville, USA) as a single-stranded DNA Ultramer® of 140bp (4nmol). We resuspended the library in nuclease-free distilled water and serially diluted it to a concentration of 50ng/uL. We used Ultramers® as templates in a PCR reaction for the formation of dsDNA fragments and library amplification. We amplified the Ultramer® DNA molecules by PCR. We performed PCR with the following program: 98°C/30 s; 25 cycles of 98°C/10 s, 60°C/15 s and 72°C/80 s; and 1 cycle of 72°C/5 min. We opted for a maximum of 25 cycles to reduce the amount of amplification biases. After amplification, we analyzed samples through gel electrophoresis to confirm that only a single band with a size around 140bp was present for each PCR reaction. After confirming the presence of single bands, we purified the samples using the Monarch® DNA gel extraction kit (NEB #T1020L). Whenever unspecific bands appeared during electrophoresis, we gel purified samples using the same kit.

**6.	Library cloning:**

We digested a total mass of 1μg of the purified library overnight (37°C, 16 hours) by HindIII-HF (NEB #R3104) and BamHI (NEB #R3136) restriction enzymes (5μL of each enzyme) in a reaction volume of 100μL. We isolated the cloning plasmid using a QIAprep spin miniprep kit (Qiagen, Germany), and digested it overnight (37°C, 16 hours) with the same restriction enzymes as the library (5uL each) in a reaction volume of 100μL. After the overnight digestion, we added 3μL of Quick CIP (M0525L) to the reaction to dephosphorylate DNA ends, and to avoid self-ligation in subsequent reactions (37°C, 2 hours). We purified samples using the Monarch® DNA gel extraction kit (NEB #T1020L).

We performed ligation following a 10:1 molar ratio of insert to vector, using 100ng of purified vector backbone, 10 units of T4 DNA ligase NEB #M0202L), and 2 µL of 10X ligation buffer (M0202L, NEB) in a 20 µL ligation reaction. We incubated the mixture at 20-22°C for ~16 h, followed by 10 min of inactivation at 65°C. We purified the ligation product using an in-house ethanol precipitation protocol (see section *DNA purification through ethanol precipitation*), resulting in 10 µL of resuspended DNA samples. Whenever recovery efficiency was low, we purified samples using the Monarch® DNA PCR/Gel Extraction Kit (NEB #T1020L), following the original protocol provided with the kit, and eluted the purified DNA in 15 µL of dH2O. We transformed *E. coli* cells (SIG10-MAX^®^) with 5 µL of the purified ligation by electroporation (see section *Electroporation*).

After transformation, we plated 50 µL of recovered cultures on LB agar plates, and incubated overnight for subsequential *cfu* (colony forming unit) counting and transformation efficiency estimation. We diluted the remaining volume of the recovered cultures (950 µL) in 9 mL of LB medium supplemented with chloramphenicol (50 µg/mL) and grew them overnight. Subsequently, we aliquoted the cultures in 1 mL cryotubes with glycerol (20% v/v final concentration) and stored them at -80°C. Based on our *cfu* counting, we estimated the mean transformation efficiency (maximal achievable library size) within this experimental setup as 10^6^ cells per transformation for the SIG10-MAX^®^strain. From the agar plates, we picked 20 colonies for colony PCR followed by Sanger sequencing to preliminarily assess library diversity (NightSeq® service, Microsynth, Switzerland).

**7.	Analysing and sorting cells**

In preparation for cell sorting, we grew cells harboring the library in LB medium supplemented with chloramphenicol. Specifically, we separately grew 1mL of frozen aliquots of transformed cells and a streak of cells harboring a control plasmid (promoterless pCAW-Sort-Seq plasmid, without sfGFP expression) overnight in a 50ml Falcon tube with 9mL of LB medium supplemented with 50 µg/mL of chloramphenicol. We then diluted the overnight cultures in LB medium supplemented with 50 µg/mL chloramphenicol in a 1:100 ratio (v/v), and grew the cultures for 5 h to late-exponential/initial-stationary phase (200RPM, 37°C). After that, we diluted 50µL of the cell cultures in 1mL of filtered cold Dulbecco’s PBS (Sigma-Aldrich #D8537) in 15 mL FACS tubes.

We performed FACS-sorting on a FACS Aria III flow cytometer (BD Biosciences, San Jose, CA) with a 70 µm nozzle for droplet formation. We used a 488 nm laser to detect forward scatter (FSC) and side scatter (SSC) with a 488nm/10nm band-pass filter. We set the flow rate to 1.0 and diluted samples if necessary to obtain a cell count of approximately 5000 events/second. Because bacterial cells are very small, we decreased the limit of particle detection (threshold) to the minimum possible (200 arbitrary units on both FSC and SSC channels). If background noise (such as caused by particles from the medium) was too high, we increased this threshold to a maximum of 500 arbitrary units on both the FSC and SSC channels. Next, we measured FSC-H and SSC-H of cells containing our negative control plasmid and set our FSC and SSC voltages to have our bacterial population on scale (centered in the software visualization panel, **Supplementary Figure S4**). We sorted and binned cells by sfGFP fluorescence (FITC channel, 488nm laser, emission filters 502LP, 530nm/30nm). We chose the FITC channel voltage such that the median fluorescence of the negative control sample (promoterless pCAW-Sort-Seq plasmid, without sfGFP expression) was between 0 and 100 (arbitrary units) on the FITC-H axis.

We set sorting gates on the FITC-H axis as follows: First, we recorded autofluorescence of the negative control cell culture. The median autofluorescence from this control served as the upper boundary of the lowest bin (B1) for the experimental population. Then we recorded the fluorescence of 10^6^ cells expressing sfGFP and harboring the library, without sorting. The choice of this number (10^6^) of cells was based on the combination of library size and the estimated loss of library diversity post-sorting (a reduction of up to 70% in sequence diversity was found in similar sort-seq studies^18^). Thus, we estimated that 975,000 cells would be required to represent the whole library (N= 65,536) with at least 5 cells harboring a copy of each sequence and accounting for diversity loss. We then proceeded to set our binning gates. We took the lower bound of the highest bin (B13) for the experimental population to correspond to the 95th percentile of the fluorescence distribution of this population. We chose boundaries between intermediate bins with equidistant spacing on a binary logarithmic (log_2_) scale. After we had the gates in this way, we calculated the fractions of the previously 10^6^ recorded cells that were inside each gate.

We determined the number of cells to be sorted into each of the 13 bins from the fraction of cells we had previously recorded in each of the bins, such that the total number of sorted cells was equal to 10^6^. We sorted cells into 1.5 mL Eppendorf tubes with 500 µL of LB medium each. We kept the tubes cooled to 4 °C to halt growth while sorting, and during the sorting of subsequent samples. We carried out the sorting procedure in three replicates derived from three independent library transformations.

We added 1mL of LB without antibiotics to each tube and removed 20 uL of the resulting volume for serial dilutions. We transferred the remaining liquid culture (980 uL) to 50 mL falcon tubes and allowed each culture to recover for 2 hours (37°C, 220 rpm). After recovery, we added 9mL of LB supplemented with chloramphenicol, and grew the cultures overnight for freezing them in glycerol stock aliquots, reassessing our binning procedure (see below), and extracting plasmid DNA for subsequent PCR and sequencing steps. We used the 20 uL initially removed from each 1mL culture for preparing two serial dilutions (10^-4^ and 10^-6^) that were plated on LB-Cm agar plates (200uL per plate) to estimate the post-sorting viability through cfu counting. By knowing how many cells were sorted into each bin, we can estimate how many cells would be expected in our dilutions and compare this number with the cfu counts we observed. In this way we estimated that on average (across bins) 73% of cells remained viable (standard deviation: 15%). We also estimated the genetic diversity of the library through Sanger sequencing of colonies (NightSeq® service, Microsynth, Switzerland).

For reassessing our binning procedure, we re-grew binned cultures (directly on the next day after sorting, using overnight recovered cultures, or by re-growing frozen aliquot stocks) and measured their expression distributions by flow cytometry. This procedure reproduced the original expression measurement (**Supplementary Figure S5**). It also allowed us to ask whether the resulting fluorescence distributions had the same geometric mean as those we had previously obtained (before sorting) for each sorting gate. The geometric mean is often preferred in this type of analysis over the arithmetic mean, because it is less affected by the presence of outliers in the data ^19,20^. It is also a more accurate representation of the central tendency of data that is log-normally distributed, which is often the case with flow cytometry fluorescence measurements^19,20^.

**8.	DNA extraction and DNA preparation for sequencing**

We diluted 500 uL of individual glycerol stocks of each replicate subpopulation of sorted cells (i.e., cells from each “bin” of fluorescent intensity) in 5mL of LB supplemented with chloramphenicol in 15mL Falcon tubes, and grew the resulting cell culture overnight (16 hours, 37°C, 220 rpm). On the next day, we isolated plasmids from each culture using a QIAprep® spin miniprep kit (Qiagen, Germany). In order to allow the sequencing of multiple pooled samples (multiplexing), we barcoded our regulatory region through PCR with specific HPLC-purified primers (**Supplementary Table S6**) provided by Eurofins (Konstanz, Germany). We added barcodes only at the 5’region of the amplicon through a PCR reaction. We performed the PCR reaction with the Q5 high-fidelity polymerase in triplicate for each sample. We calculated the primer melting temperatures (Tm) following the NEB Tm calculator (<https://tmcalculator.neb.com/#!/main>) for a primer concentration of 500nM. We performed PCR with the following program: 98°C/30 s; 25 cycles of 98°C/10 s, 64°C/30 s and 72°C/30 s; and 1 cycle of 72°C/2 min.

After PCR amplification, we digested the reaction products with both the restriction enzyme DpnI (NEB #R0176L) and Exonuclease I (NEB #M0293L) in order to remove traces of genomic DNA, plasmids, and single-stranded DNA that could interfere with sequencing. The Master Mix we used for a single digestion harbored 1µL of 10x CutSmart® Buffer (NEB #B6004S), 1µL of Exonuclease I (NEB #M0293L), 1µL of DpnI restriction enzyme (NEB #R0176L), and 7µL of distilled nuclease-free water. For each PCR product, we added 10µL of the Master Mix. We incubated the reaction for 1 hour at 37°C, following 15 minutes at 80°C for deactivation of the enzymes.

After digestion, we purified samples using the Monarch® DNA PCR/Gel Extraction Kit (NEB #T1020L. We also analyzed samples through gel electrophoresis to confirm that only a single band with a size around 150bp was present for each of them. After confirming the expected bands, we pooled the samples for the different bins of each replicate equimolarly to a total mass of 2,600 ng and a volume of 100 µL (26ng/µL of DNA) in 1.5mL Eppendorf tubes. We then sent the pooled samples for adapter ligation and sequencing at Eurofins (NGSelect Amplicons® on Illumina HiSeq), obtaining 15 million paired-end reads (2 × 150 bp) for all samples.

**9.	Data analysis:**

*9.1	Filtering and preparing sequencing reads*

We processed the sequencing data using a combination of in-house Python and awk scripts, as well as standard bioinformatics tools. Briefly, we first trimmed sequences to remove Illumina adaptors, merged the paired reads and separated them into different files according to their sequence barcodes, which reflect the specific fluorescence bins from which they originated, **Supplementary Table S6**.

In order to remove Illumina adaptor sequences from the paired-end reads (in both forward and reverse orientations), we used Cutadapt ^21^ with the following options:

^136136^

*cutadapt -j 8 -e 0.1 --no-indels --overlap=8 --discard-untrimmed \*

*-a "^\$FWD...\$REV_RC;max_error_rate=0.2;min_overlap=6" \*

*-A "^\$REV...\$FWD_RC;max_error_rate=0.2;min_overlap=6" --pair-filter=any \*

*-o 'Sample_${sample}_trimmed_1.fastq.gz' \*

*-p 'Sample_${sample}_trimmed_2.fastq.gz' \*

*--max-ee=2 -l=114 \*

*$reads*

The “read1” adapter we used was:

5’-AGATCGGAAGAGCACACGTCTGAACTCCAGTCA-3’

The “read2” Adapter we used was:

5’-AGATCGGAAGAGCGTCGTGTAGGGAAAGAGTGT-3’

Because our amplicons were short (153 bp) and the paired-reads thus fully overlap, we had to merge them. We performed this process with the software FLASH^22^. After trimming and merging the paired-end reads, we performed the demultiplexing process using the following FLASH options:

*flash -t $ncores $reads -O -m 60 -M 140 -z -o 'Sample_${sample}_merged'*

Subsequently, we used the open-source tool FastX (<http://hannonlab.cshl.edu/fastx_toolkit/>) to identify and retain only high quality sequences (quality threshold of Q = 33).

After these steps we filtered our data further, retaining only sequences that showed neither mutations nor indels in the *sfgfp* regulatory region outside the variable TFBS library region. For this purpose, we developed an in-house python script. Subsequently, we used in-house *awk* and R ^23^ scripts to compile the data as a single table containing all unique sequences as rows, and the number of reads for each bin as columns. We used this table to calculate the average gene expression level driven by each sequence.

*9.2	Calculating repression levels:*

In sort-seq experiments, sequences can appear in more than a single fluorescence bin due to both random mis-sorting^24,25^ and stochastic gene expression noise^26^. Therefore, following previous studies^27–29^, we estimated expression levels for each TFBS sequence as the weighted average of bins in which the sequence was observed. Specifically, we multiplied, for each sequence, the number of times (*x_i_*) that we had observed the sequence in each bin *i* by an integer representing that bin (*w_i_*=1,2,3,4,…,13) and averaged these values by the total number of read counts for that sequence. In mathematical terms, we calculated the weighted average

e=$\sum_{i=1}^{n} \frac{(x_{i}*w_{i})}{\sum_{i=1}^{n} x_{i}}$

to quantify the expression level e driven by the sequence.

This procedure resulted in a continuous distribution of expression levels e within the interval (1, 13). Since high expression values represent high GFP expression and, therefore, low-affinity binding of TetR to a TFBS variant, we reversed the scale to facilitate biological interpretation, where high values correspond to strong repression and low GFP expression. Specifically, we calculated

$$r = e_{max} + 1 - e$$

as a measure of the repression level r conferred by a sequence, where e_max_=13 is the maximum possible value for the expression level. In addition, to obtain a metric that lends itself to more intuitive interpretation, we normalized the previously calculated average repression levels, dividing them by the wild type repression level. Hence, the repression scores we used in this study range from 0 to 1.25, where low values represent TetR TFBS variants that bind TetR with low-affinity, and high values represent variants that bind TetR with high-affinity and repress gene expression strongly. The value of one corresponds to the repression conferred by the “wild-type” *tetO2* binding site*.*

*9.3	Combining the data from triplicates:*

Firstly, we removed from further analysis all sequence variants that were not present in all triplicates. Secondly, we eliminated all sequences that did not have a minimum of 30 reads in all 13 bins (**Supplementary Figure S8**). We then calculated the repression levels (see Calculating individual average expression levels and repression levels section) for each remaining sequence in each replicate. Thirdly, we calculated the coefficient of variation of repression levels for each sequence (across replicates) as a measure of how widely distributed the repression levels were among replicates. We filtered out sequences with a coefficient of variation above 0.5. In doing so, we followed common practice in transcriptional studies using fluorescent reporters, which accept a coefficient of variation up to this magnitude as admissible due to transcriptional noise and technical variability of measurements ^26^. Lastly, we combined our datasets by averaging across replicates the repression levels conveyed by each remaining sequence.

*9.4	Frequency matrix and sequence logo:*

We generated frequency matrices of nucleotides for TetR TFBSs from each fluorescence bin by counting the frequency of each nucleotide at each variable position of the *tetO2* library. We generated heatmaps representing the frequency matrices and DNA sequence logos in R. A sequence logo consists of a stack of letters at each position of a DNA sequence, where the relative sizes of the letters indicate their frequency in the sequence. The total height of the stack corresponds to the information content of that position, in bits ^30,31^.

Such logos are graphical representations of informational properties of DNA. When mutated, characters with high information content are more likely to lead to a loss of binding (repression) than characters with low information content.

*9.5	Creation of genotype networks and network metrics:*

We used in-house Python script and the python package *igraph*^32^ to generate directed genotype networks. These are graphs in which TetR TFBS variants are nodes (vertices, genotypes), and variants that differ in a single nucleotide are connected by an edge. Each edge is directed, i.e., it corresponds to a binding-score-increasing mutation and points from the variant with lower repression level to the neighbor with higher repression level. Each vertex of this network is associated with the corresponding DNA sequence and the associated repression level. We extracted the largest weakly connected subgraph (giant component^33^) of the network and used it for all further analyses. This giant component comprises the vast majority (97%) of sequenced genotypes. We used in-house Python and R scripts for all network analyses described in the sections below.

*Epistasis:*

Epistasis, non-additive interactions between mutations, can impose severe constraints on molecular evolution, because the mutations that are beneficial in one genetic background may be deleterious in another^34^. Epistasis can be classified as magnitude, simple sign, or reciprocal sign epistasis, depending on the sign (i.e., positive or negative) of the fitness effect of individual mutations and their combinations^34^ —with increasingly detrimental effects on landscape navigability^35,36^. In magnitude epistasis, the effect of a mutation on repression varies depending on the genetic background but the sign of this effect (increasing or decreasing repression) does not. Simple sign epistasis occurs if one single mutant has a lower repression level than both the wild type and the double mutant, while the other single mutant has a repression level that is intermediate to the wild type and double mutant. Reciprocal sign epistasis occurs when both mutations independently decrease repression but their combination increases repression. Thus, the presence of reciprocal sign epistasis is a necessary condition for the existence of multiple peaks in an adaptive landscape ^35,36^.

To conduct epistasis calculations, we employed a method that involved identifying all "squares" in the genotype network using the *motifs* function from the *igraph* library in R. Each square consisted of a "wild-type" sequence, a double-nucleotide mutant, and the corresponding two single mutants (see **Supplementary Figure S17** for more information on epistasis squares). We assessed epistasis for each square along a single axis by selecting the highest-repression sequence as the double mutant. Our analysis categorized the landscape into three groups: no sign epistasis, simple sign epistasis, and reciprocal sign epistasis, which we explained in detail in **Supplementary Table S1** and **Supplementary Figure S17**. The no sign epistasis category included both magnitude epistasis and additivity (no epistasis), without differentiating between them, as neither affected peak accessibility^11,108^. Finally, we determined the proportion of all squares whose constituent mutations interacted based on these classifications (see **Supplementary Table S1** and **Supplementary Figure S17**).

*Peaks:* A peak is a genotype (TetR binding site variant) whose neighbors all convey lower repression than itself. Two peaks are connected if they are neighbors and convey the same repression.

We refer to the genotype with the highest repression level as the summit or global peak^33,37^.

*Accessible Paths:* A mutational path through the genotype network is accessible, if and only if the repression level increases for each mutational step along the path^33,37^. We enumerated accessible paths of all lengths (mutational steps) exhaustively.

*Basins of attraction:* The basin of attraction of a peak comprises all TetR binding site variants from which accessible paths to the peak exist. We refer to the basin’s size as the number of variants in the basin. We determined basin sizes by exhaustive enumeration.

*Overlap between basins.* The basins of attraction of different peaks may comprise overlapping sets of variants. To determine the overlap between two basins *B_1_* and *B_2_*, we used the Jaccard index *J*^38,39^, which is equal to the size of the intersection between two sets of variants divided by the size of their union:

$$J=\frac{B1\cap B2}{B1\cup B2}$$

*9.6	Principal component analysis*

In order to investigate the proximity of peaks and other variants in sequence space, we prepared our data with a one-hot encoding method. A one-hot encoding represents each categorical value (nucleotide at each position of a DNA string) as a binary vector of length 4.

It thus converts an entire DNA string of length L into a 4xL binary matrix. We used the binary matrix to perform the PCA analysis. We performed the PCA analysis using the R base function *prcomp()*.

**10.	Simulated adaptive walks**

We simulated the adaptive evolution of a population on the adaptive landscape by performing three different types of random walks, a “greedy adaptive” random walk, a “uniform adaptive” random walk, and a random walk based on Kimura’s model of fixation probability.

For all simulations, we assumed that only point mutations occur and that the time it takes for a point mutation to become fixed in a population is much shorter than the time it takes for a new mutation to appear that will eventually go to fixation. This scenario is also referred to as the strong selection weak mutation (SSWM) scenario ^28,40–42^. Under this scenario, evolving populations are monomorphic most of the time, i.e., all individuals have the same genotype. This scenario is realistic when the product of effective population size *N* and mutation rate *μ* is small (*Nμ* < 1), which is the case for ***E. coli*** (N=1.8×10^8^, μ=2×10^-10^) ^43^. In this scenario, adaptive evolution on a fitness landscape can be modelled as an adaptive random walk.

In the greedy adaptive random walk, we assume that only the mutation that conveys the largest fitness advantage is fixed in a population. We initiated one greedy random walk from each non-peak genotype and terminated the walk once no genotype with higher fitness was reachable, i.e., a fitness peak was reached. Because every greedy random walk is deterministic in the absence of neutral mutations, it is sufficient to initiate one greedy random walk per starting genotype.

With the uniform adaptive random walk we relax the conditions of the greedy random walk by allowing all fitness-increasing mutations to fix with equal probability. We randomly chose 1,000 non-peak starting genotypes and simulated 1000 uniform random walks for each of these starting genotypes. Random walks terminate when they reach a fitness peak.

Both types of adaptive walks ignore the possibility of genetic drift, which allows mutations with no or negative fitness effects to become fixed in a population. A convenient model for random walks that permit genetic drift uses fixation probabilities computed by Kimura^44–46^ i.e., *f_ij_* = (1 – *e*^-2^*^s^*) / ( 1 – *e*^-2^*^Ns^* ), where *f_ij_* is the probability of fixing mutation *j* in the background of genotype *i*, *N* is the effective population size, and *s* is the selection coefficient, i.e. the difference in fitness between genotypes *i* and *j*^44,45^. For a given pair of genotypes, the only parameter of this model is the effective population size *N*, for which we explored values of *N*=10^8^, *N*=10^5^ and *N*=10^2^. Generally, the smaller the population size, the larger the probability that neutral or deleterious mutations become fixed in a population.

For these “Kimura” adaptive walks, we chose 1000 random starting genotypes for each of the three values of *N*, and simulated 1000 random walks for each of them. At each step, we randomly picked a mutation *j*, generated a random number on the interval [0, 1], and considered the mutation to become fixed if the random number fell within the interval [0, *f_ij_*]. Computationally, we accelerated this process by precomputing all fixation probabilities for all genetic backgrounds, and then used the *Python* function numpy.choice to generate a random sample from the multinomial distribution of fixation probabilities at each step^47^. Because of genetic drift, Kimura random walks do not necessarily terminate when they reach a fitness peak. For this reason, we simulated each such random walk for 1,000 mutational steps, which allows multiple fitness peaks to be visited.

**Supplementary Tables**

| **Table S1. Network metrics** | |
| --- | --- |
| Number of nodes (genotypes) | 17,765 |
| Number of peaks | 2,092 |
| Number of low peaks^1^ | 2,034 (97.2%) |
| Number of high peaks^2^ | 58 (2.8%) |
| Number of squares^3^ | 83,100 |
| Magnitude epistasis or additivity^4,5^ | 35% |
| Simple sign epistasis^5^ | 34% |
| Reciprocal sign epistasis^5^ | 30% |

^1^ Low peaks are peaks with repression levels below the wild sequence*.* Percentages refer to the proportion of all genotpyes.

^2^ High peaks are peaks with repression levels above the wild-type sequence. Percentages refer to the proportion of all peaks.

^3^ A square represents the connection between a focal sequence (ab) and a double mutant (AB) via two single mutants (Ab and aB).

^4^ This category includes both magnitude epistasis and additivity (no epistasis) without distinguishing them, because neither of the two subcategories affects peak accessibility^11,108^

^5^ Percentages refer to the proportion of all squares.

**Table S2. Table of strains**

| **Strain** | **Genotype** |
| --- | --- |
| SIG10-MAX from Sigma Aldrich | F- mcrA Δ(mrr-hsdRMS-mcrBC) endA1 recA1 Φ80dlacZΔM15 ΔlacX74 araD139 Δ(ara,leu)7697galU galK rpsL nupG λ- tonA (StrR) |

**Table S3. Table of plasmids used in this study**

| **Name** | **Selective antibiotics (concentration µg/ml)** | **Relevant features** | **Source** | **T, °C** | **Description** |
| --- | --- | --- | --- | --- | --- |
| **pCAW-Sort-Seq** | Chloramphenicol (50) | pBBR1, TetR, *sfgfp* | This study | 37 | Vector used for library generation and sort-seq |
| **pCAW-Sort-Seq-Neg** | Chloramphenicol (50) | pBBR1, TetR, promoterless *sfgfp* | This study | 37 | pCAW-Sort-Seq vector without a promoter for *sfgp* (negative control) |

**Table S4. Primers for engineering the pCAW-sort-seq plasmid**

| **Name** | **Sequence** | **Function** |
| --- | --- | --- |
| pCAW_frag1_F | CGTCCGACTTACGGAAGGTAGATtttacggc | Linearizing the pCAW-Sort-Seq fragment1 for Gibson Assembly |
| pCAW_frag1_R | CTCGTGCCTAACGGAAGGTAGATtttacggc | Linearizing the pCAW-Sort-Seq fragment1 for Gibson Assembly |
| pCAW_frag2_F | TAAGATTGCCACGGAAGGTAGATtttacggc | Linearizing the pCAW-Sort-Seq fragment2 for Gibson Assembly |
| pCAW_frag2_R | AGGCCTGACTACGGAAGGTAGATtttacggc | Linearizing the pCAW-Sort-Seq fragment2 for Gibson Assembly |

**Table S5 Primers for constructing libraries**

| **Name** | **Sequence** | **Function** |
| --- | --- | --- |
| Ultramer_ds_F | TTCTCAAAAGCTTCCTGCAGTATTC | Amplifying Ultramer® libraries |
| Ultramer_ds_R | cggaaagcacatccggtgac | Amplifying Ultramer® libraries |
| TFBS_R | CCGTTTGTAGCATCACCTTC | Sequencing the TFBS region |
| pCAW_Gibs_Lib_F | gtctgatgagtccgtgaggacg | Linearizing the pCAW-Sort-Seq plasmid |
| pCAW_Gibs_Lib_R | GAGAAAAGAAAACCGCCGATCCTG | Linearizing the pCAW-Sort-Seq plasmid |
| Ultramer_Gibs_F | GGTGGACAGGATCGGCGGTTTTCTTTTCTCTTCTCAAAAGCTTCCTGCAGTATTC | Amplifying Ultramer® libraries for Gibson Assembly |
| Ultramer_Gibs_R | ggctgtttcgtcctcacggactcatcagaccggaaagcacatccggtg | Amplifying Ultramer® libraries for Gibson Assembly |

**Table S6. Primers with barcodes for demultiplexing sequencing bins:**

| **Name** | **Sequence** |
| --- | --- |
| Bin_1_F | **AGTCTCGGCA**ACGGAAGGTAGATtttacggc |
| Bin_2_F | **GATATAGCTC**ACGGAAGGTAGATtttacggc |
| Bin_3_F | **CGTCCGACTT**ACGGAAGGTAGATtttacggc |
| Bin_4_F | **CTCGTGCCTA**ACGGAAGGTAGATtttacggc |
| Bin_5_F | **TAAGATTGCC**ACGGAAGGTAGATtttacggc |
| Bin_6_F | **AGGCCTGACT**ACGGAAGGTAGATtttacggc |
| Bin_7_F | **GTCAATCTTC**ACGGAAGGTAGATtttacggc |
| Bin_8_F | **ATGACGGTAA**ACGGAAGGTAGATtttacggc |
| Bin_9_F | **AGGCTCAAGG**ACGGAAGGTAGATtttacggc |
| Bin_10_F | **GCTCAGTAAT**ACGGAAGGTAGATtttacggc |
| Bin_11_F | **ACGATGAAGT**ACGGAAGGTAGATtttacggc |
| Bin_12_F | **GAGCAGATAT**ACGGAAGGTAGATtttacggc |
| Bin_13_F | **CGATAGCGAG**ACGGAAGGTAGATtttacggc |
| Bin_R_1 | tcctcacggactcatcagac |

Barcodes are represented in bold letters

**Supplementary figures:**

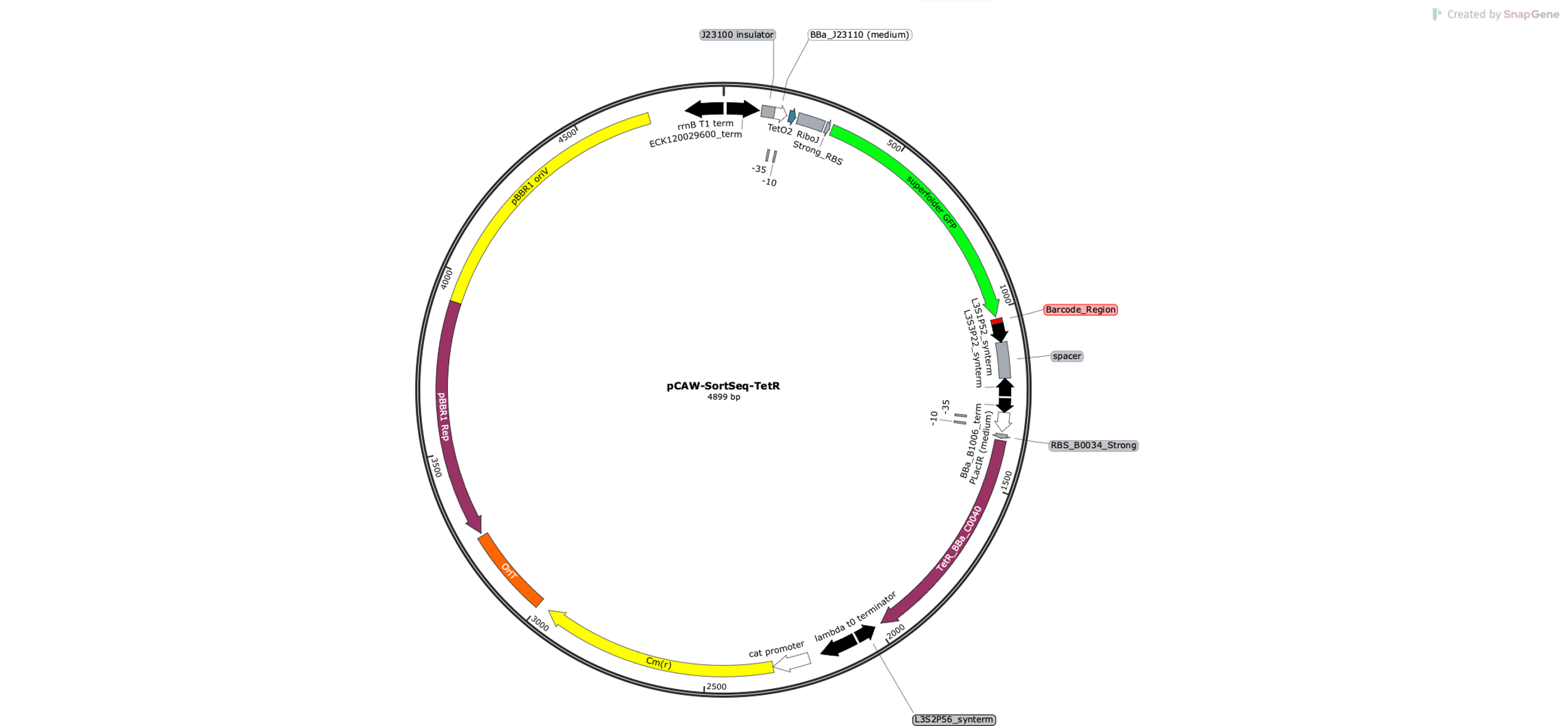

**Supplementary Figure S1. The pCAW-SortSeq-TetR plasmid.** The plasmid system pCAW-SortSeq encodes a broad-host, low-copy number replication origin (pBBR1 replication origin – 5 to 10 copies) ^70^, an interchangeable regulatory region where the TFBS is located and placed between a constitutive promoter (BBa_J23110 ^71^), a superfolder GFP (*sfgfp*) fluorescent reporter gene^72^, as well as a *tetr* gene. The *tetr* gene is derived from the original Tn10 transposon^51,73^ under the control of a low-strength constitutive promoter (pLac promoter variant developed by ^74^ ).

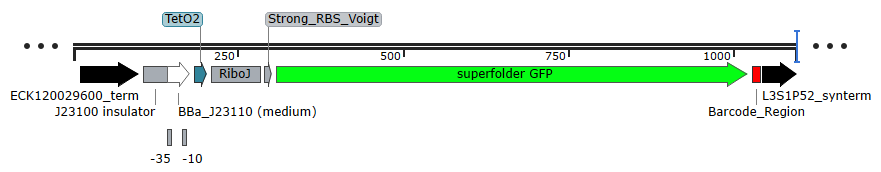
 **Supplementary Figure S2. The repression measurement module of the pCAW-SortSeq-TetR plasmid.** This module encodes the promoter insulator *J23100_Insulator* upstream of the *BBa_J23110* constitutive promoter from ^87^. Promoter insulators are transcriptional terminators that alleviate contextual effects of upstream sequences on promoter regions. Bold underlined letters represent -35 and -10 boxes. The constitutive promoter of this module is the *BBa_J23110* promoter from the iGEM registry of parts (<http://parts.igem.org/Part:BBa_J23110>) ^71^, which has medium promoter strength. We placed the *tetO2* sequence^66^ at the +10 position relative to the *sfgfp* transcription start site, which is the optimal distance for synthetic repression^85^. The transcriptional insulator RiboJ has been described in ^79^. It is a synthetic ribozyme that removes 5’UTR interferences with variable TFBS sequences in the mRNA by self-cleavage. We obtained the strong synthetic RBS sequence from ref. ^93^, the reporter gene superfolder GFP (*sfgfp)* from ref. ^72^, and the strong synthetic transcriptional terminator (*synterm*) from ref. ^89^**.**

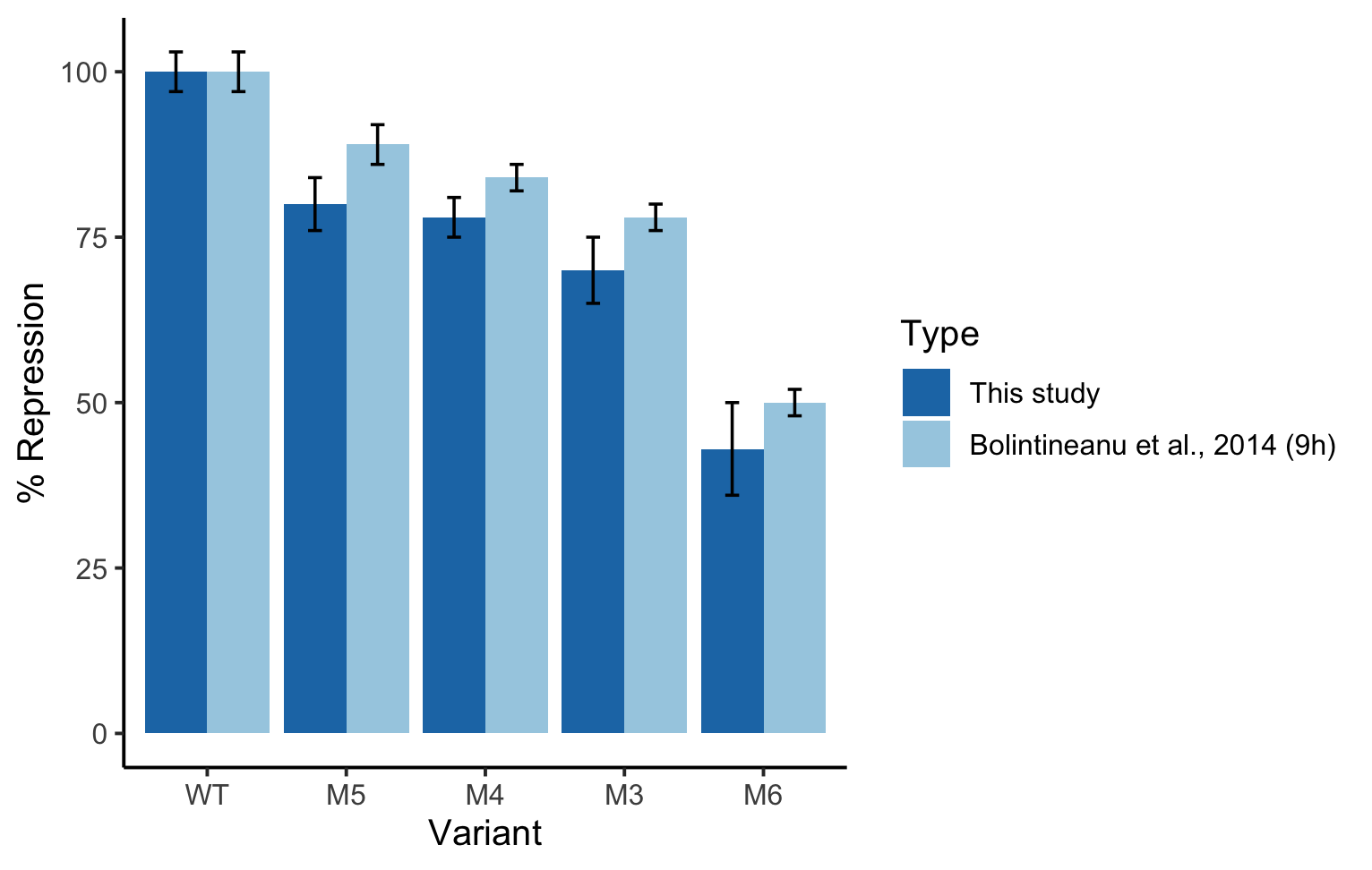

**Supplementary Figure S3. Validating the pCAW-SortSeq-TetR plasmid with *tetO2* mutants.** We validated the pCAW-SortSeq-TetR plasmid with *tetO2* mutants by measuring the percentage of repression (vertical axis) for each of five *tetO2* variants (WT, M5, M4, M3, M6, horizontal axis). For calculating percentages of repression, we measured GFP fluorescence distributions for each variant in triplicate in a flow cytometer, divided the mean fluorescence (over triplicate measurements) of each variant by the mean of the WT *tetO2*, and multiplied by 100. Light blue bars represent the data from a previous study^17^ characterizing each of the five TetR TFBS variants. Dark blue bars correspond to measurements obtained in the present study. Error bars represent standard deviations among replicates.

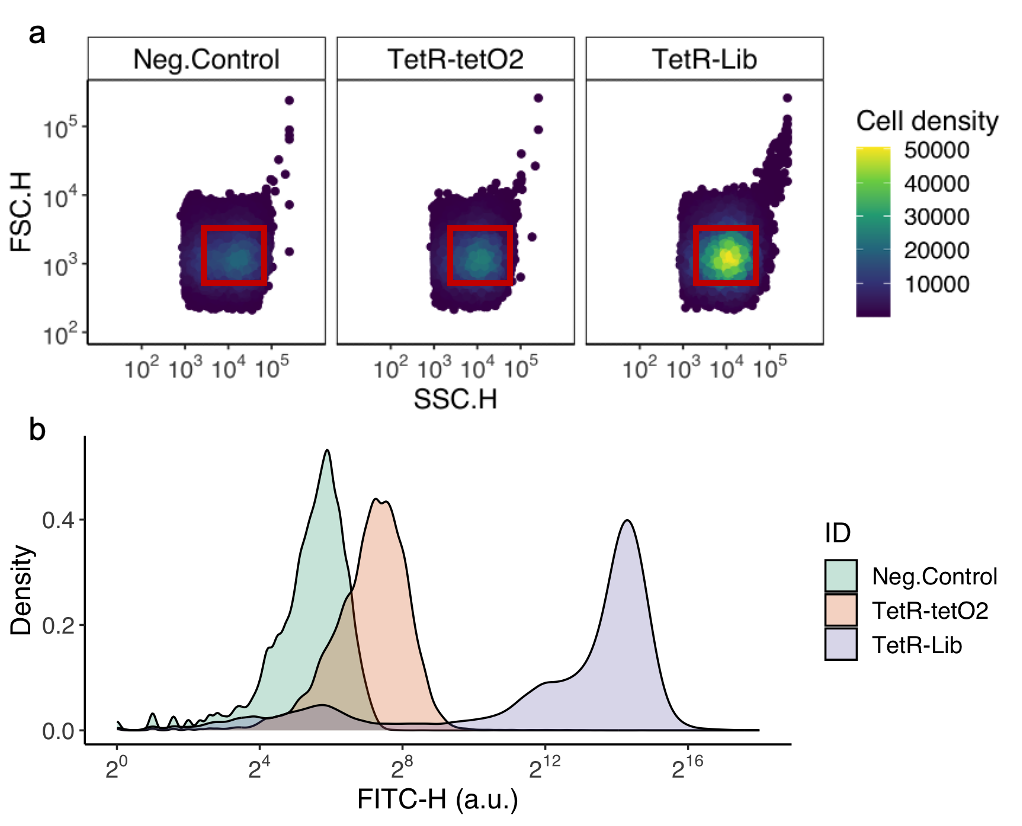

**Supplementary Figure S4. Flow cytometry gating and fluorescence expression levels for controls and library. a. Gating of three representative cell populations.** We measured the forward (FSC.H, vertical axis) and side scatter (SSC.H, horizontal axis) for 200,000 cells per sample. Inside each grid, we depict individual cells as circles. Heatmap colors represent population densities (see color legend). From left to right: cells harbouring a negative control plasmid (pCAW with promoterless GFP), a positive *tetO2* control (pCAW with the wild-type *tetO2* instead of the mutant library), and *tetO2* variants (pCAW with variant library). All populations were grown in the absence of anhydrotetracycline. The red box represents the region of each scatter plot where cell density was the highest, from which cells were sorted in subsequent experiments. **b.** **Fluorescence distributions of three cell populations transformed with different plasmids.** Density plot with the fluorescence distribution for the same three samples described above (see color legend). The horizontal axis represents the range of values for GFP fluorescence as FITC-H (arbitrary units, note the log_2_ scale). The vertical axis represents the relative frequency of observations for each fluorescence value on the horizontal axis. Density smoothing was performed using a Gaussian kernel function to create a smooth density plot – *ggplot2* package^48^.

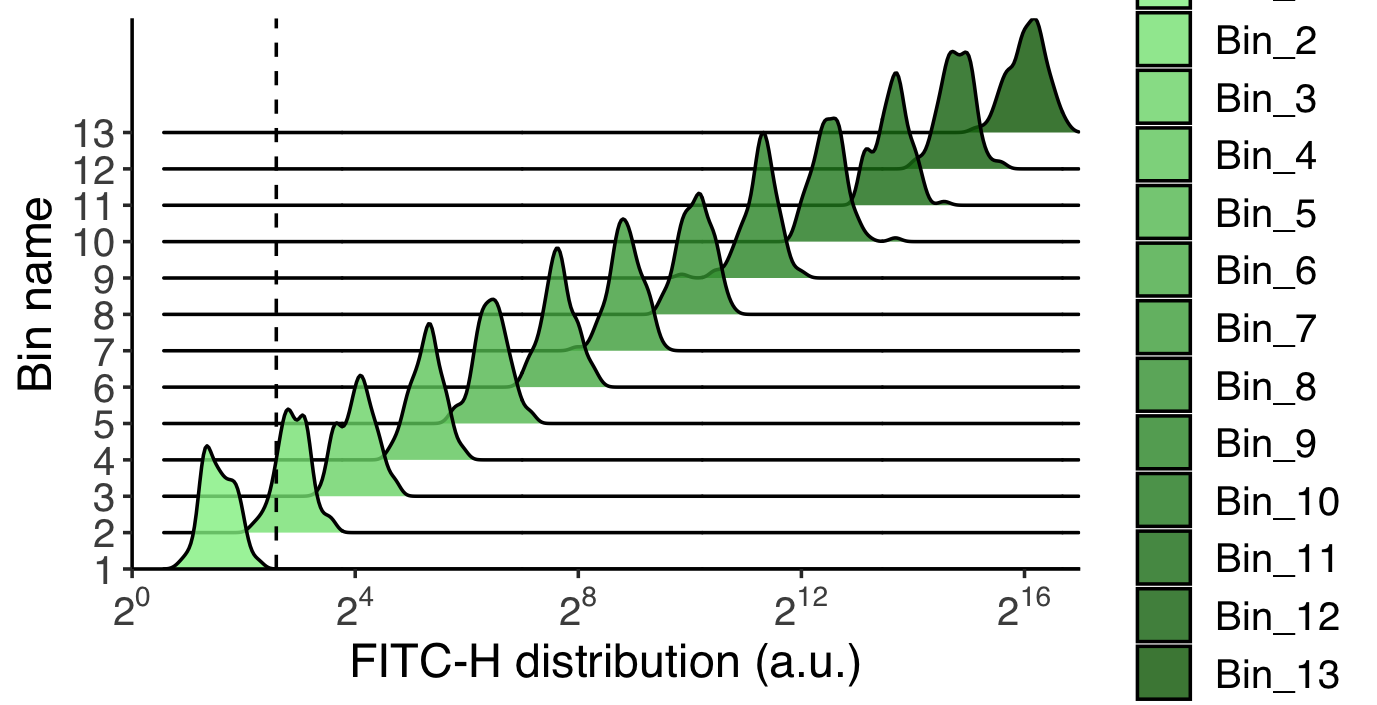

**Supplementary Figure S5. Distribution of fluorescence levels after sorting of cells expressing the library into fluorescence bins.** The distribution of fluorescence values for each bin is shown as individual density plots. The color gradient represents changes in GFP expression levels (as quantified in the FITC-H channel, arbitrary units) across the horizontal axis, with lighter green corresponding to low GFP expression (and thus higher repression) and darker green corresponding to higher GFP expression (and thus weaker repression). The vertical dashed line represents the autofluorescence threshold based on the obtained geometric mean calculated over the fluorescence distribution for the negative control population. We determined the distribution of values for each bin from a population of 200,000 cells.

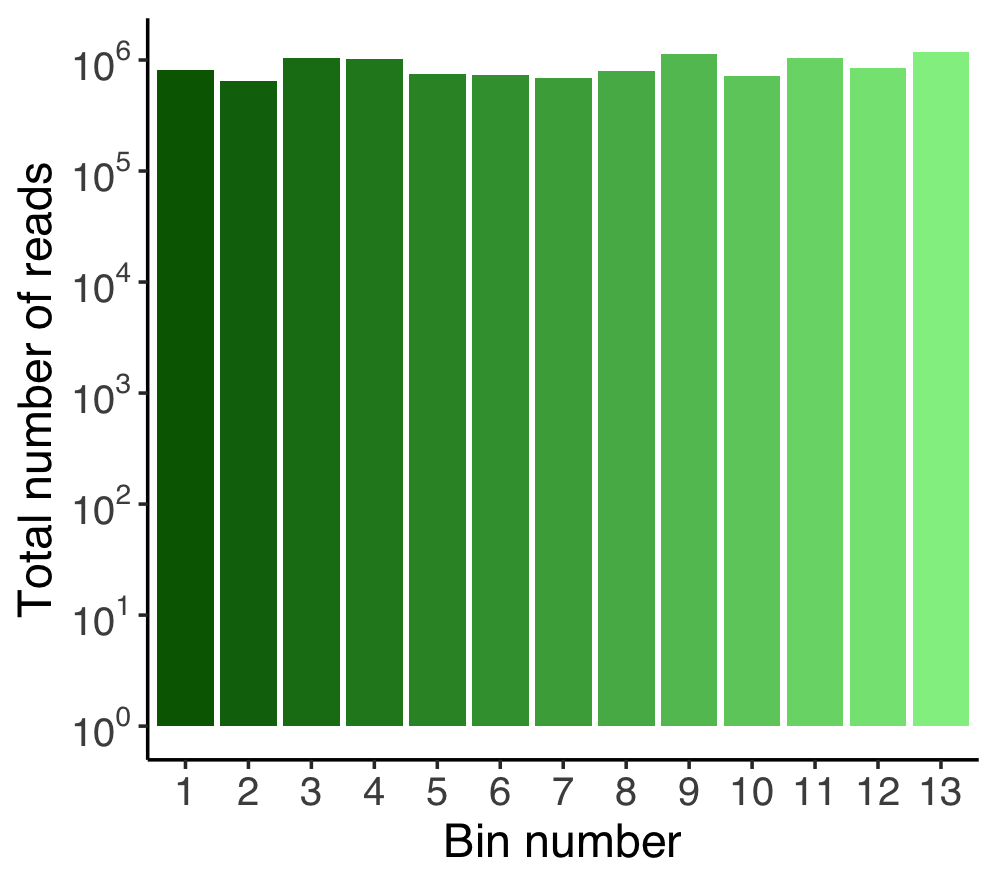

**Supplementary Figure S6. Distribution of sequence reads per bins**. Gradient bar colours depict GFP expression levels across bins. The total number of sequence reads in each bin is represented on the vertical axis on a logarithmic scale.

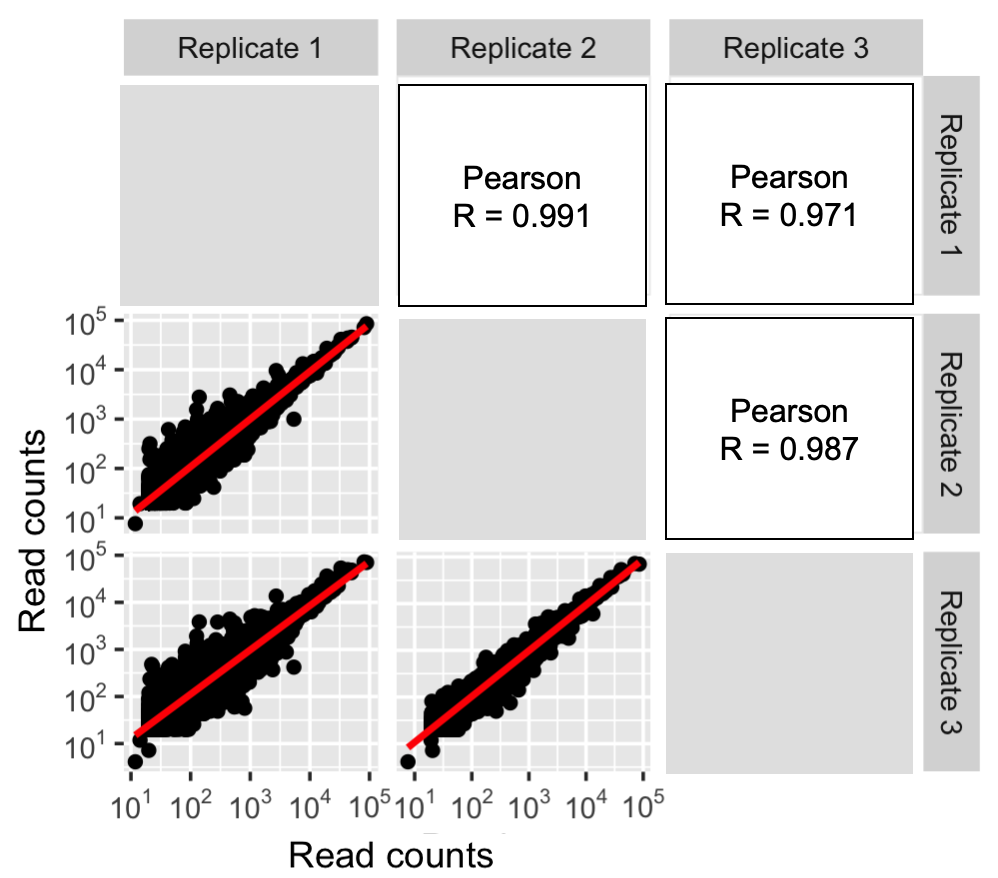

**Supplementary Figure S7. Correlation of read coverage among replicates of the *tetO2* mutant library**. Note the logarithmic scale in all panels. Correlation plots are represented as scatter plots in the lower panels, the red line in each plot is the x=y line. The R in upper figure panels represent the Pearson correlation coefficients calculated for each pair of replicates.

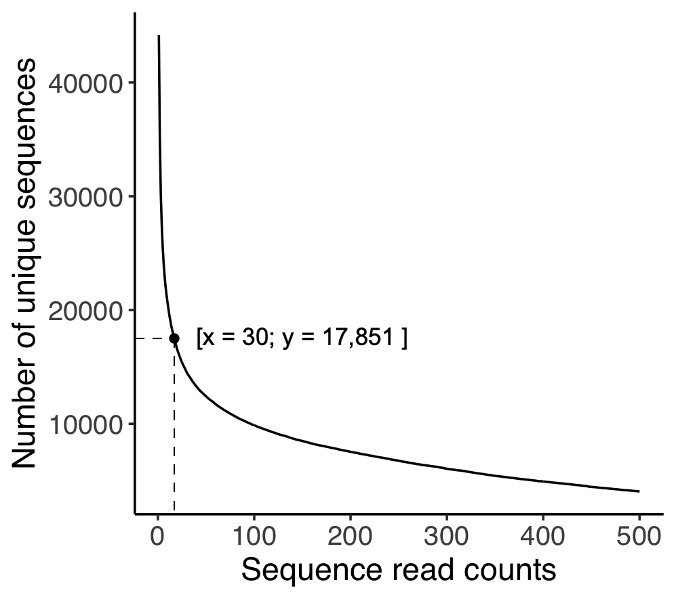

**Supplementary Figure S8. Number of genotypes obtained by choosing different sequencing depth (read count) cutoffs.** The number of sequence reads represents different read count thresholds in the interval (1,500), i.e., the minimum number of required counts to include a genotype in our analysis. The dashed line shows the number (17,851) of unique sequences that pass the threshold of 30 reads that we used in our analysis.

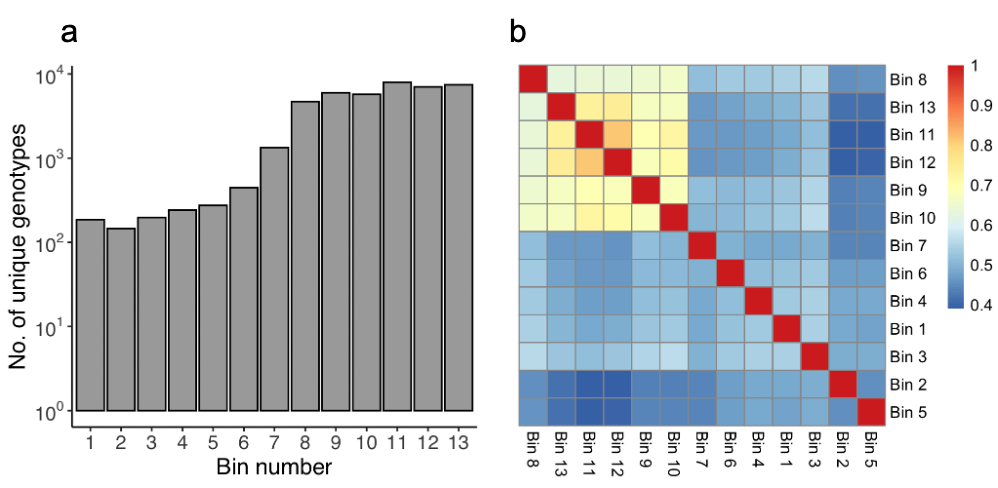

**Supplementary Figure S9. Genotypes per fluorescence bin and overlaps between them. a. Number of individual genotypes per bin.** The vertical axis shows the number of unique genotypes per bin for each of the 13 fluorescence bins (horizontal axis) into which we sorted cells. Low fluorescence bins correspond to TFBS variants conferring strong repression, of which there are fewer, hence there are also fewer unique genotypes in these bins. **b.** **Heatmap of the fraction of genotypes shared between different bins.** The data is represented as a symmetric matrix of pairwise fractional overlaps calculated using the Jaccard index coefficient ^38,39^ between all possible pairs of the13 bins. Each row and each column corresponds to a bin. Red values represent complete overlap between bin sequences and dark blue values represent the minimum overlap observed (40%). We ordered and clustered bins using and Euclidean distance with a complete-linkage clustering method in which the Euclidean distance measures the similarity or dissimilarity between bins, and the complete-linkage clustering merge clusters based on the distance between their farthest points. Note that overlaps do not consider read counts. For example, two bins might share 40% of their sequences but the read count between the same sequence in each bin might differ by orders of magnitude. Note also that the overlap is greatest for high fluorescence bins, which also contain the most genotypes (panel a). The higher genotype overlap between higher bins (Bins 8-13) can be explained by a higher number of cells sorted into these bins in comparison to lower bins.

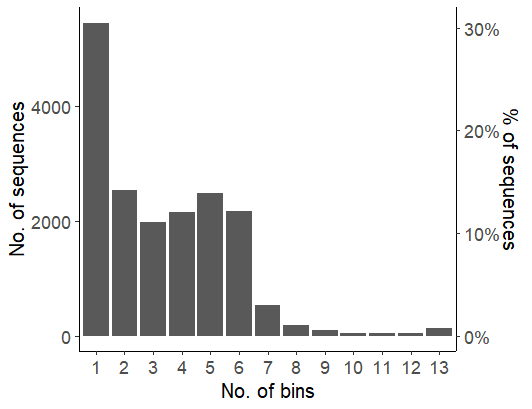

**Supplementary Figure S10. Number of bins per genotype.** The figure shows a histogram of the distribution of the number of bins (horizontal axis) into which each TFBS variant was sorted. The secondary vertical axis on the right shows the same information, but as a percentage of the total number of sequences (100% =17,851 sequences). 32% of sequences were sorted only into a single bin; 65% of sequences were sorted into 2 to 6 bins.

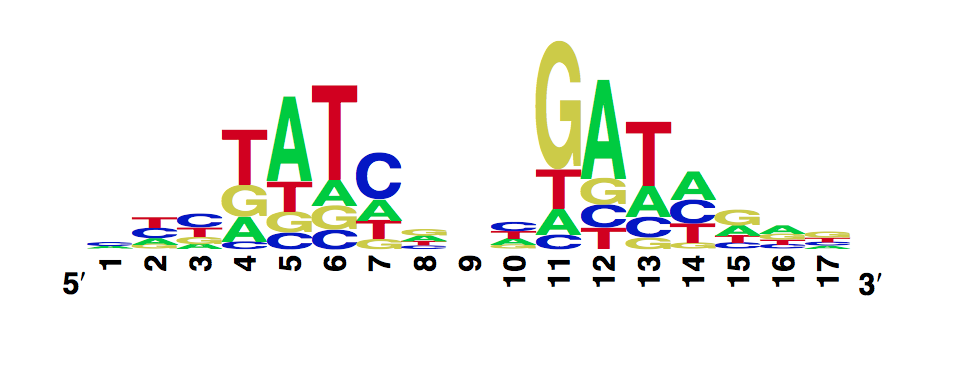

**Supplementary Figure S11. DNA Sequence logo obtained by a previous study.** Using the MITOMI^49^ in vitro technique, the 2011 iGEM team of the École polytechnique fédérale de Lausanne (EPFL) studied the DNA binding landscape of the wild-type TetR sequence. To do so, it designed and generated a library of double-stranded DNA sequences that cover all possible single base substitutions within the *tetO2* binding site sequence. Based on that library the team measured dissociation constants of each variant relative to the average constant of all the *tetO2*-like variants of the library. Then, it determined the specificity of TetR for the binding site variant sequences, expressed as a position-weight matrix (PWM). The figure shows the corresponding DNA sequence logo. Original figure available at <https://2011.igem.org/Team:EPF-Lausanne/Our_Project/TetR_mutants/MITOMI_data>).

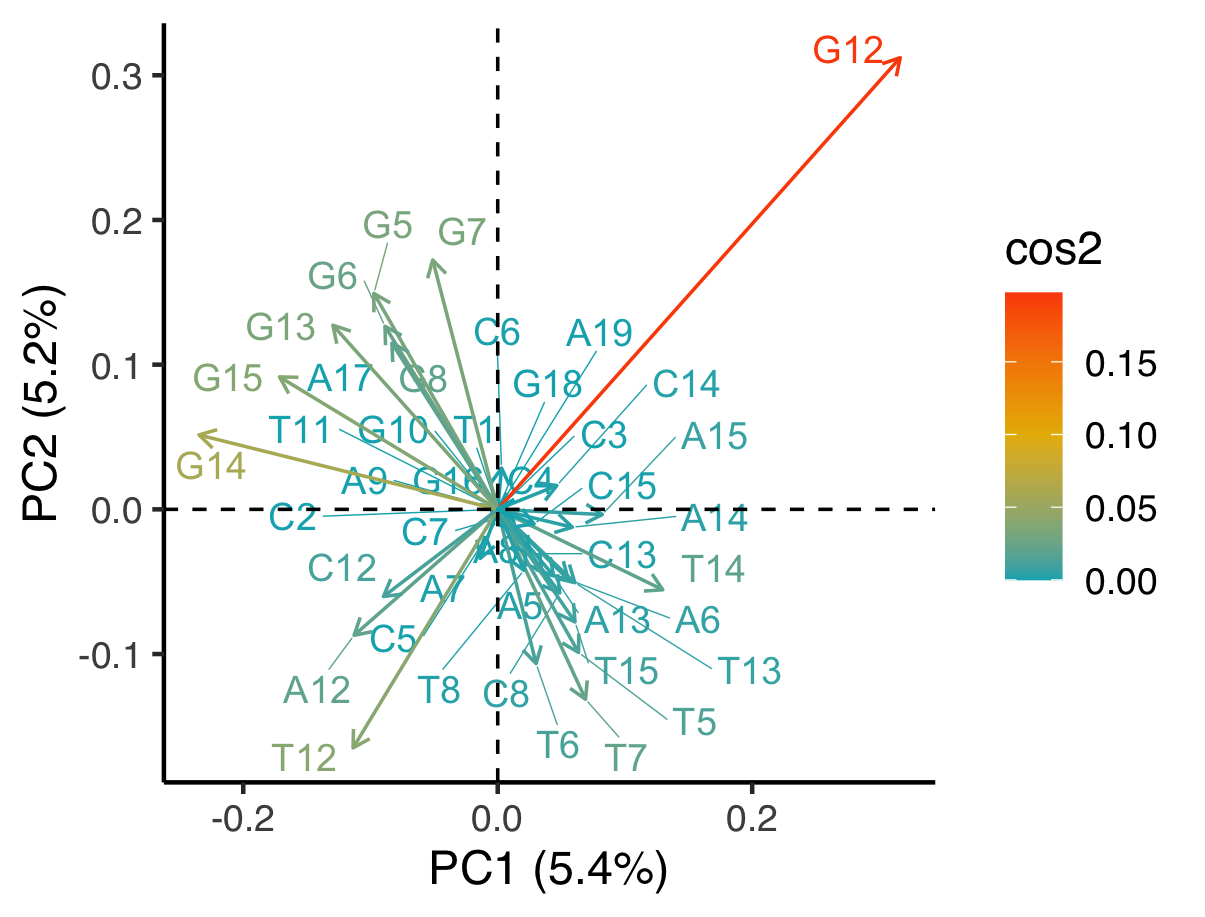

**Supplementary Figure S12. The contribution of individual *tetO2* nucleotides in principal component analysis.** A PCA (Principal Component Analysis) contribution plot is a way to visualize the relative importance of different variables to the variation observed in the data. In this plot, the contribution of each variable is expressed as the square of the cosine of the angle (cos2) between the variable's vector (column representing the presence/absence of a base letter at each position in the sequence) and each principal component axis. This quantity is represented as an arrow that indicates the correlation of the variable with PC1 and PC2, the two principal components that capture the largest amount of variation in the data. Both length and color of the arrow represent the contribution of the variable to the variation observed in the data. A high cos2 value (red) indicates that the variable is strongly correlated with the principal component, and therefore makes a large contribution to the variation observed in the data. A low cos2 value (blue) indicates that the variable is weakly correlated with the principal component, and therefore makes a small contribution to the variation observed in the data. The arrow size represents the importance of the variable's contribution relative to other variables in the plot. Each nucleotide is represented with as a base letter (A, T, C, G) followed by a number that indicates the position of that base in the binding site sequence (e.g. G12 stands for a guanine at position 12 of the binding site).

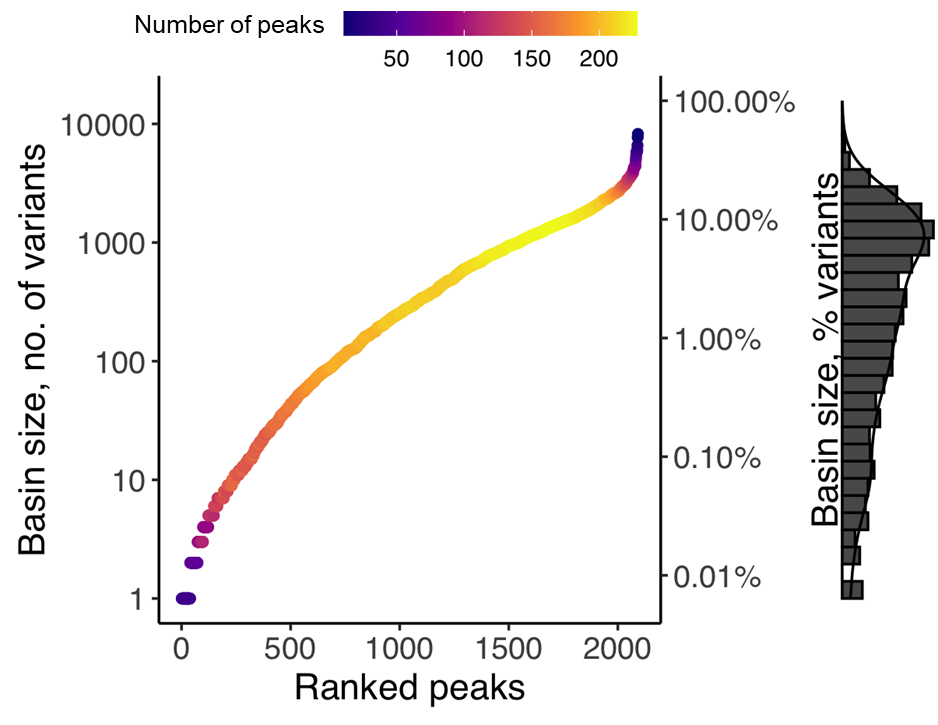

**Supplementary Figure S13. The distribution of basin sizes among peaks.** Peaks are ranked along the horizontal axis according to the size of their basins of attraction (vertical axis). The secondary vertical axis on the right represents basin size as a percentage of variants (100% =17,765 variants). Heatmap colors represent the number of peaks at each position of the ranked scatterplot (see color legend). The marginal histogram on the right shows the distribution of basin sizes.

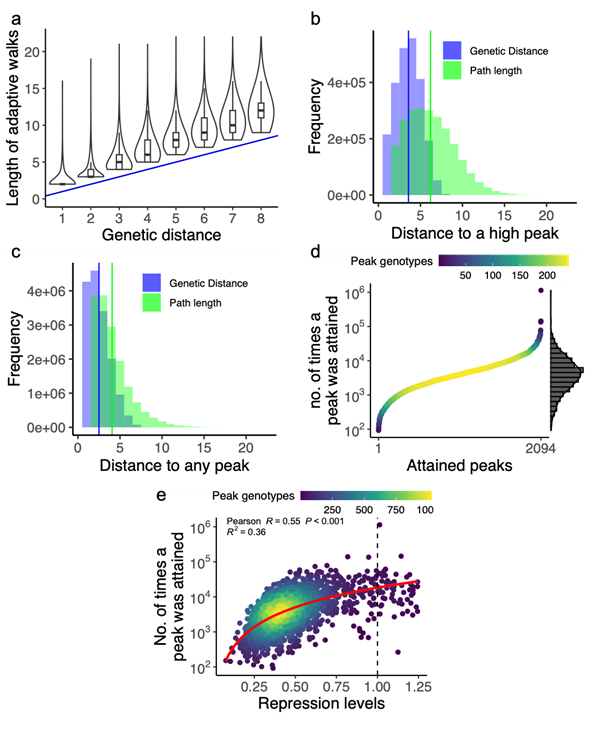

**Supplementary Figure S14. Adaptive walks in which each mutational step is chosen with uniform probability among all repression-increasing steps.** Data displayed here are based on 1,000 adaptive walks starting from each non-peak genotypic variant (N= 15.671). **a. Adaptive walks leading to high repression peaks are predominantly short.** The vertical axis presents the number of mutational steps in adaptive walks that initiate from a random variant and converge at a high repression peak. The horizontal axis reflects the shortest genetic distances between the starting variant and the attained peak. The blue line (y=x) signifies the most direct distance to a high repression peak, equated by the genetic distance. Violin plots summarize the shape of distributions with a probability density function. A wider probability density function indicates that a value on the y-axis occurs more frequently, and a narrower density function indicates that a value occurs less frequently. Each box covers the range between the first and third quartiles (IQR). The horizontal line within the box represents the median value, and whiskers span 1.5 times the IQR. Values beyond the 1.5 IQR interval are shown. Adaptive walks were only marginally longer than shortest paths. **b.** **Accessible paths to high repression peaks tend to be short.** The blue histogram shows the distribution of the genetic distances for all pairs of variants and their respective attainable high peaks. The green histogram shows the distribution of the number of mutational steps for shortest accessible paths between variants and their respective attainable peaks. **c.** **Most accessible paths to** **any (high or low)** **repression peak are short.** The blue histogram shows the distribution of the genetic distances for all pairs of variants and their respective attainable peaks. The green histogram shows the distribution of the number of mutational steps for shortest accessible paths between variants and their respective attainable peaks (high or low). **d. Some peaks are attained more frequently than others.** We ranked all peaks along the horizontal axis according to the number of times they are reached across all adaptive walk simulations (N= 15.671 × 10^3^, vertical axis, note logarithmic scale). The marginal density histogram on the right shows the distribution of the number of times each peak was reached from an individual variant. Heatmap colors represent the number of peaks at each position of the ranked scatterplot (see color legend). **e. High peaks tend to be attained more often.** The scatter plot shows the repression level conveyed by a peak variant (horizontal axis) and the number of times a peak with this repression level was reached across all adaptive walks (N= 15.671 × 10^3^, vertical axis, note logarithmic scale). The dashed vertical line represents the repression level for the wild-type sequence. The red curve represents a semi-logarithmic linear regression line for the data, and the grey shade around it represents its 95% percent confidence interval. *R* is the linear Pearson correlation coefficient, and *R^2^* is the goodness of fit of the logarithmic linear regression model (N= 15.671). Heatmap colors represent the number of peaks at each position of the ranked scatterplot (see color legend).

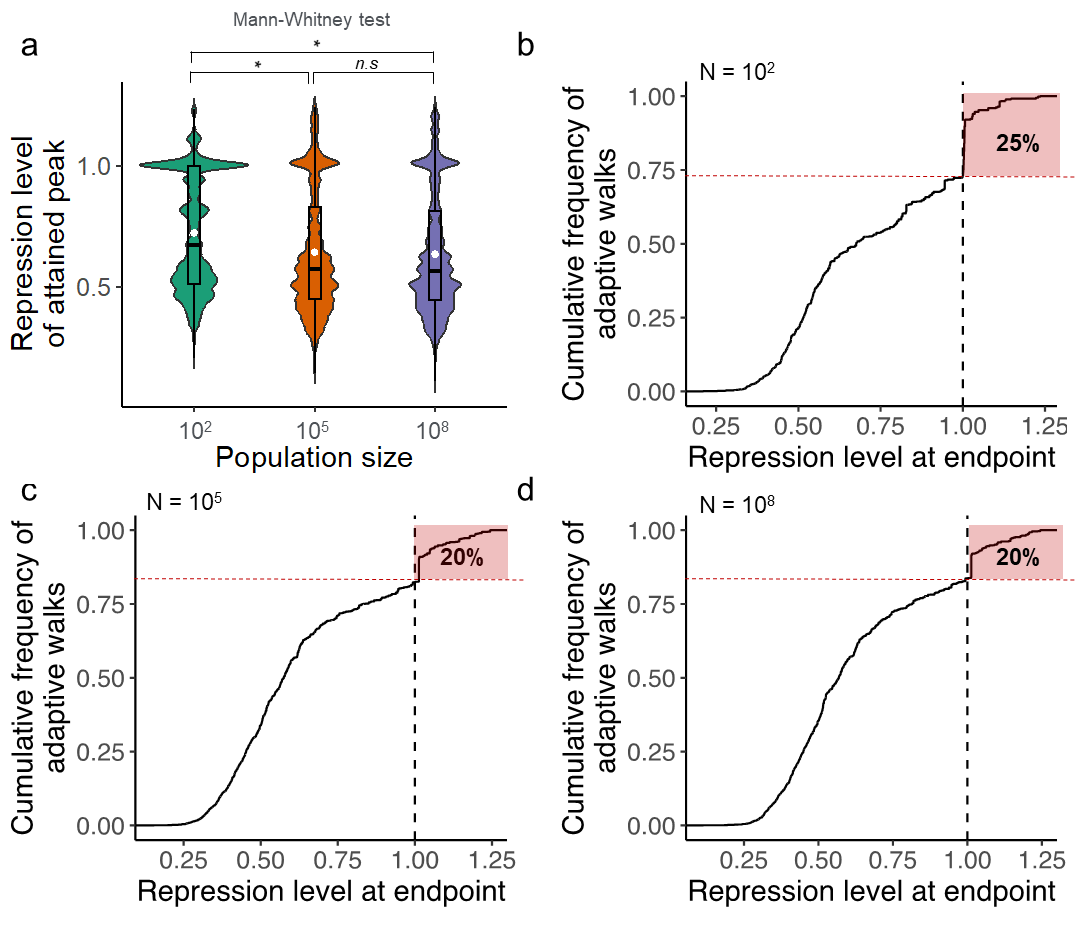

**Supplementary Figure S15. Adaptive walks using the Kimura model with different population sizes**. Data displayed here are based on 10^6^ adaptive walks starting from 10^3^ random variants (10^3^ random walks per variant) for small (N=10^2^), medium (N=10^5^) and large (N=10^8^) population sizes. **a. Distribution of repression levels of attained peaks for each population size.** Violin plots summarize the shape of distributions for each population size (horizontal axis). Each box covers the range between the first and third quartiles (IQR). The horizontal line within the box represents the median value, and whiskers span 1.5 times the IQR.. The white circle inside each boxplot represents the mean of each distribution, which is 0.72±0.34, 0.64±0.25 and 0.63±0.25 (mean ±s.d.) for small, medium, and large population sizes, respectively. The median repression level of attained peaks for small populations (10^2^) is significantly higher than that for medium and large populations (two-sided Mann–Whitney U = 425,000,000, n1 = 10^6^, n2 = 10^6^ , p = 2.13 × 10^-15^). **b-d.** **Small population sizes attain higher peaks.** Each panel shows the cumulative distribution of repression values reached by 10^6^ adaptive walks starting from 1,000 random variants (1,000 walks per variant) in the landscape (Supplementary methods). The population size of each panel is represented by the *N* letter on the upper left of each plot. The dashed vertical line *x*=1 shows the repression value of the wild type. The area highlighted in red corresponds to the percentages of adaptive walks that reached peaks with repression 1 or greater; 25%, 20% and 20% of adaptive walks for small, medium, and large population sizes, respectively.

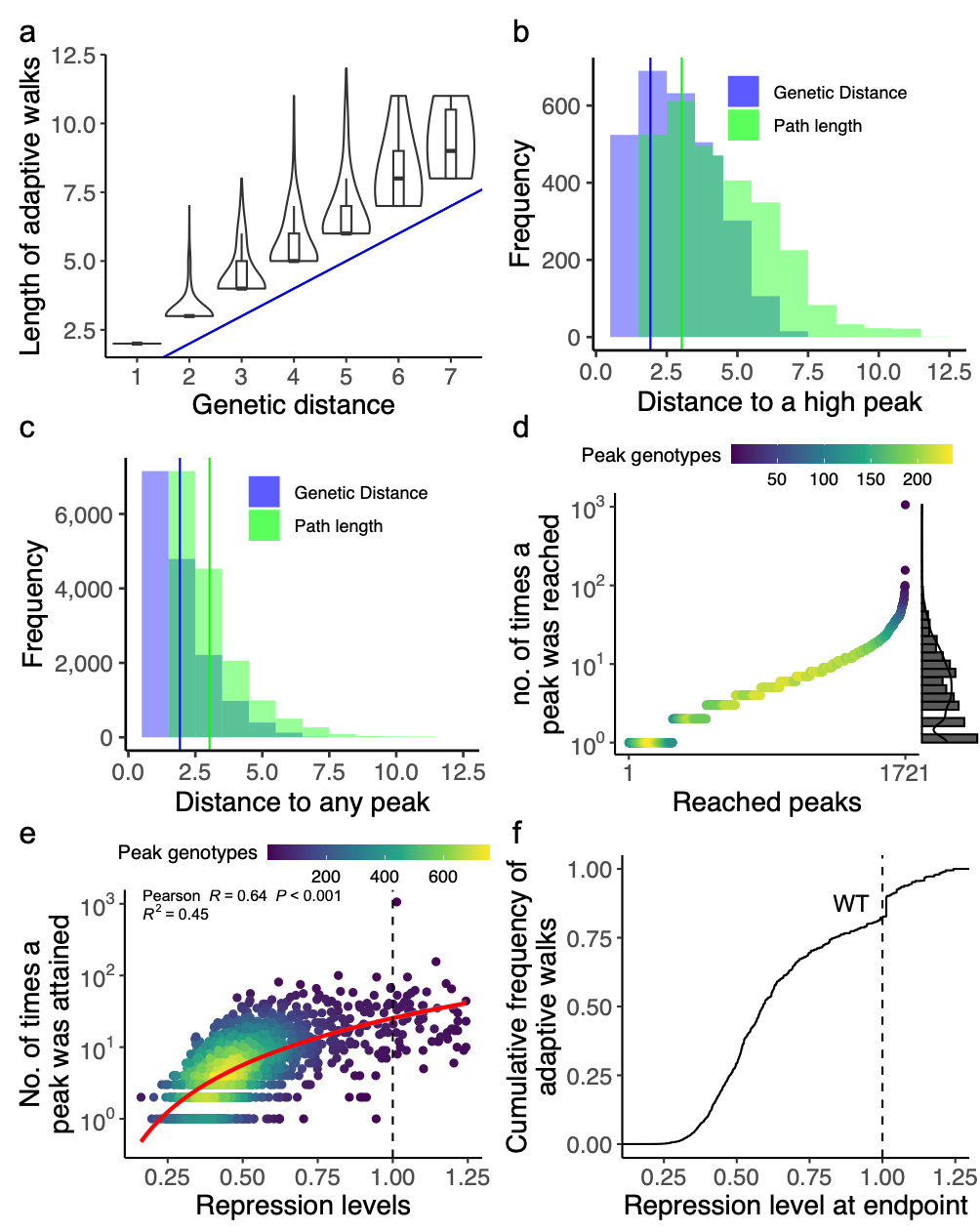

**Supplementary Figure S16. Greedy adaptive walk simulations.** We performed one greedy adaptive walk starting from each of N= 15,671 non-peak genotypes (Supplementary Methods 10). Each such walk is deterministic**. a. Adaptive walks leading to high repression peaks are predominantly short.** The vertical axis presents the number of mutational steps in adaptive walks that initiate from a random variant and converge at a high repression peak. The horizontal axis reflects the shortest genetic distances between the starting variant and the attained peak. The blue line (y=x) signifies the most direct distance to a high repression peak, equated by the genetic distance. Violin plots summarize the shape of distributions with a probability density function. A wider probability density function indicates that a value on the y-axis occurs more frequently, and a narrower density function indicates that a value occurs less frequently. Each box covers the range between the first and third quartiles (IQR). The horizontal line within the box represents the median value, and whiskers span 1.5 times the IQR. Values beyond the 1.5 IQR interval are shown. Adaptive walks were only marginally longer than shortest paths. **b.** **Accessible paths to high repression peaks tend to be short.** The blue histogram shows the distribution of the genetic distances for all pairs of variants and their respective attainable peaks. The green histogram shows the distribution of the number of mutational steps for shortest accessible paths between variants and their respective attainable peaks. **c.** **Most accessible paths to** **any (high or low)** **repression peak are short.** The blue histogram shows the distribution of the genetic distances for all pairs of variants and their respective attainable peaks. The green histogram shows the distribution of the number of mutational steps for shortest accessible paths between variants and their respective attainable peaks (high or low). **d. Some peaks are attained more frequently than others.** We ranked all peaks along the horizontal axis according to the number of times they are reached across all adaptive walk simulations (N= 15.671, vertical axis, note logarithmic scale). The marginal density histogram on the right shows the distribution of the number of times each peak was reached from an individual variant. Heatmap colors represent the number of peaks at each position of the ranked scatterplot (see color legend). **e. High peaks tend to be reached more often.** The scatter plot shows the repression level conveyed by a peak variant (horizontal axis) and the number of times a peak with this repression level was reached across all adaptive walks (N= 15.671, vertical axis, note logarithmic scale). The dashed vertical line represents the repression level for the wild-type sequence. The red curve represents a semi-logarithmic linear regression line for the data, and the grey shade around it represents its 95% percent confidence interval. *R* is the linear Pearson correlation coefficient, and *R^2^* is the goodness of fit of the logarithmic linear regression model (N= 15.671). Heatmap colors represent the number of peaks at each position of the ranked scatterplot (see color legend). **f. High repression peaks are attainable through adaptive evolution.** The panel shows the cumulative distribution of repression values reached by 10^3^ adaptive walks starting from each non-peak variant in the landscape (Methods). The dashed vertical line at *x*=1 shows the repression value of the wild type. Only 20% of adaptive walks reached a repression value of 1 or higher.

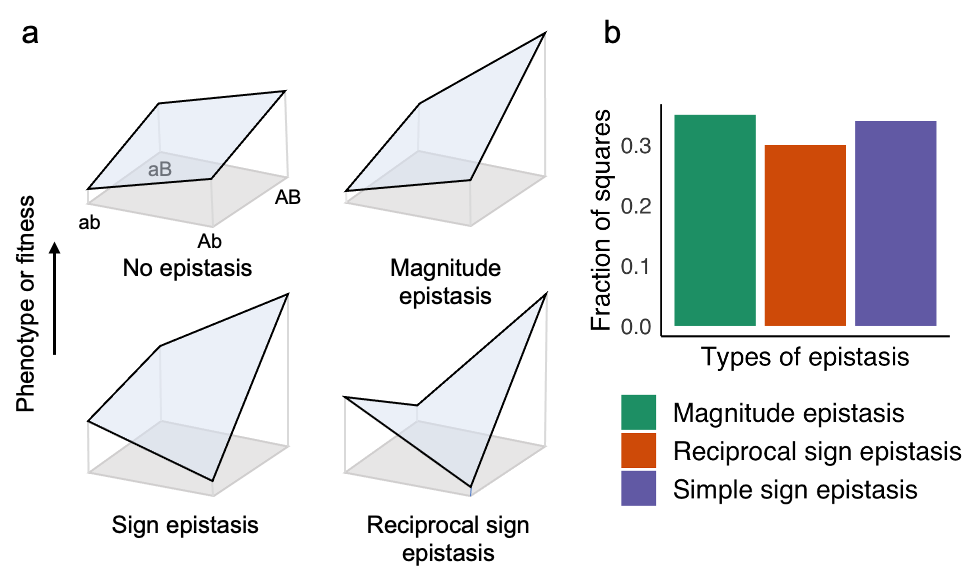
 **Supplementary Figure S17. Epistatic interactions can influence adaptation and hence ruggedness of adaptive landscapes.** **a. Overview of epistatic interactions.** A ‘wild-type’ sequence (ab) can change to a double mutant (AB) via the single mutations Ab or aB. The upper left panel shows a mutational path without epistasis, where the repression value of the double mutant is the sum of the repression contributions of both single mutants (additive interaction). Magnitude epistasis changes the magnitude, but not the sign of a resulting repression value. In the example, the repression value associated with the AB genotype is higher than the sum of the repression values for Ab and aB (second panel). Sign epistasis occurs when one single mutant (Ab) has a lower repression value than both the ‘wild type’ and the double mutant, while the other single mutant (aB) shows intermediate repression (third panel). In reciprocal sign epistasis, both single mutations decrease repression individually, but increase repression jointly (in the double mutant). Note that the relationship between fitness and repression is dependent on the studied system. In the case of TetR, we have assumed that fitness and repression are positively associated, based on previous studies exploring such relationship^50–53^ Figure adapted from^54^. **b. Prevalence of three types of epistasis in the landscape.** The bar plot shows the prevalence of the three major types of epistatic interactions (panel a) in our data (*N*=83,100 double mutant pairs). The category called "no sign epistasis" comprises of both magnitude epistasis and additivity (no epistasis), without any differentiation between them, as neither of them have an impact on peak accessibility ^55,56^.

REFERENCES

1.	Warren, D. J. Preparation of highly efficient electrocompetent Escherichia coli using glycerol/mannitol density step centrifugation. *Anal Biochem* **413**, 206–207 (2011).

2.	Silva-Rocha, R. *et al.* The Standard European Vector Architecture (SEVA): a coherent platform for the analysis and deployment of complex prokaryotic phenotypes. *Nucleic Acids Res* **41**, D666-75 (2013).

3.	Jahn, M., Vorpahl, C., Hübschmann, T., Harms, H. & Müller, S. Copy number variability of expression plasmids determined by cell sorting and droplet digital PCR. *Microb Cell Fact* **15**, 211 (2016).

4.	Lutz, R. & Bujard, H. Independent and tight regulation of transcriptional units in Escherichia coli via the LacR/O, the TetR/O and AraC/I1-I2 regulatory elements. *Nucleic Acids Res* **25**, 1203–10 (1997).

5.	Pédelacq, J. D., Cabantous, S., Tran, T., Terwilliger, T. C. & Waldo, G. S. Engineering and characterization of a superfolder green fluorescent protein. *Nat Biotechnol* **24**, 79–88 (2006).

6.	Meyer, A. J., Segall-Shapiro, T. H., Glassey, E., Zhang, J. & Voigt, C. A. Escherichia coli “Marionette” strains with 12 highly optimized small-molecule sensors. *Nature Chemical Biology 2018 15:2* **15**, 196–204 (2018).

7.	Kelly, J. R. *et al.* Measuring the activity of BioBrick promoters using an in vivo reference standard. *J Biol Eng* **3**, 4 (2009).

8.	Zaslaver, A. *et al.* A comprehensive library of fluorescent transcriptional reporters for Escherichia coli. *Nat Methods* **3**, 623–628 (2006).

9.	Gibson, D. G. Programming biological operating systems: genome design, assembly and activation. *Nat Methods* **11**, 521–6 (2014).

10.	Lou, C., Stanton, B., Chen, Y. J., Munsky, B. & Voigt, C. A. Ribozyme-based insulator parts buffer synthetic circuits from genetic context. *Nat Biotechnol* (2012) doi:10.1038/nbt.2401.

11.	Carr, S. B., Beal, J. & Densmore, D. M. Reducing DNA context dependence in bacterial promoters. *PLoS One* (2017) doi:10.1371/journal.pone.0176013.

12.	Yeung, E. *et al.* Biophysical Constraints Arising from Compositional Context in Synthetic Gene Networks. *Cell Syst* **5**, 11-24.e12 (2017).

13.	Chen, Y. J. *et al.* Characterization of 582 natural and synthetic terminators and quantification of their design constraints. *Nat Methods* (2013) doi:10.1038/nmeth.2515.

14.	Urtecho, G., Tripp, A. D., Insigne, K. D., Kim, H. & Kosuri, S. Systematic Dissection of Sequence Elements Controlling σ70 Promoters Using a Genomically Encoded Multiplexed Reporter Assay in Escherichia coli. *Biochemistry* **58**, 1539–1551 (2019).

15.	Jack, B. R. *et al.* Predicting the Genetic Stability of Engineered DNA Sequences with the EFM Calculator. *ACS Synth Biol* **4**, 939–943 (2015).

16.	Gibson, D. G. *et al.* Enzymatic assembly of DNA molecules up to several hundred kilobases. *Nat Methods* **6**, 343–5 (2009).

17.	Bolintineanu, D. S. *et al.* Investigation of changes in tetracycline repressor binding upon mutations in the tetracycline operator. *J Chem Eng Data* **59**, 3167–3176 (2014).

18.	Kinney, J. B., Murugan, A., Callan, C. G. & Cox, E. C. Using deep sequencing to characterize the biophysical mechanism of a transcriptional regulatory sequence. *Proceedings of the National Academy of Sciences* **107**, 9158–9163 (2010).

19.	Beal, J. Biochemical complexity drives log-normal variation in genetic expression. *Engineering Biology* **1**, 55–60 (2017).

20.	Beal, J. *et al.* Reproducibility of fluorescent expression from engineered biological constructs in E. coli. *PLoS One* **11**, (2016).

21.	Martin, M. Cutadapt removes adapter sequences from high-throughput sequencing reads. *EMBnet J* **17**, 10–12 (2011).

22.	Magoč, T. & Salzberg, S. L. FLASH: fast length adjustment of short reads to improve genome assemblies. *Bioinformatics* **27**, 2957–2963 (2011).

23.	Bunn, A. & Korpela, M. R: A language and environment for statistical computing. (2013) doi:10.1016/j.dendro.2008.01.002.

24.	Peterman, N. & Levine, E. Sort-seq under the hood: Implications of design choices on large-scale characterization of sequence-function relations. *BMC Genomics* **17**, 1–17 (2016).

25.	Trippe, B. L. *et al.* Randomized gates eliminate bias in sort-seq assays. *Protein Science* **31**, e4401 (2022).

26.	Elowitz, M. B., Levine, A. J., Siggia, E. D. & Swain, P. S. Stochastic Gene Expression in a Single Cell. *Science (1979)* **297**, 1183–6 (2007).

27.	de Boer, C. G. *et al.* Deciphering eukaryotic gene-regulatory logic with 100 million random promoters. *Nature Biotechnology 2019 38:1* **38**, 56–65 (2019).

28.	Vaishnav, E. D. *et al.* The evolution, evolvability and engineering of gene regulatory DNA. *Nature 2022 603:7901* **603**, 455–463 (2022).

29.	Lagator, M. *et al.* Predicting bacterial promoter function and evolution from random sequences. *Elife* **11**, (2022).

30.	Schneider, T. D. & Stephens, R. M. Sequence logos: a new way to display consensus sequences. *Nucleic Acids Res* **18**, 6097 (1990).

31.	Stormo, G. D. DNA binding sites: Representation and discovery. *Bioinformatics* vol. 16 16–23 Preprint at https://doi.org/10.1093/bioinformatics/16.1.16 (2000).

32.	Csardi, G. The Igraph Software Package for Complex Network Research. (2014).

33.	Aguilar-Rodríguez, J., Payne, J. L. & Wagner, A. A thousand empirical adaptive landscapes and their navigability. *Nat Ecol Evol* **1**, 0045 (2017).

34.	Bank, C. Epistasis and Adaptation on Fitness Landscapes. *https://doi.org/10.1146/annurev-ecolsys-102320-112153* **53**, 457–479 (2022).

35.	Saona, R., Kondrashov, F. A. & Khudiakova, K. A. Relation Between the Number of Peaks and the Number of Reciprocal Sign Epistatic Interactions. *Bull Math Biol* **84**, (2022).

36.	Poelwijk, F. J., Tǎnase-Nicola, S., Kiviet, D. J. & Tans, S. J. Reciprocal sign epistasis is a necessary condition for multi-peaked fitness landscapes. *J Theor Biol* **272**, 141–144 (2011).

37.	Khalid, F. *et al.* Genonets server-a web server for the construction, analysis and visualization of genotype networks. *Nucleic Acids Res* **44**, W70–W76 (2016).

38.	Jaccard, P. THE DISTRIBUTION OF THE FLORA IN THE ALPINE ZONE.1. *New Phytologist* **11**, 37–50 (1912).

39.	Papkou, A., Garcia-Pastor, L., Escudero, J. A. & Wagner, A. A rugged yet easily navigable fitness landscape of antibiotic resistance. *bioRxiv* 2023.02.27.530293 (2023) doi:10.1101/2023.02.27.530293.

40.	Bank, C., Matuszewski, S., Hietpas, R. T. & Jensen, J. D. On the (un)predictability of a large intragenic fitness landscape. *Proc Natl Acad Sci U S A* **113**, 14085–14090 (2016).

41.	Orr, H. A. The population genetics of adaptation: the adaptation of DNA sequences. *Evolution* **56**, 1317–1330 (2002).

42.	Gillespie, J. H. MOLECULAR EVOLUTION OVER THE MUTATIONAL LANDSCAPE. *Evolution (N Y)* **38**, 1116–1129 (1984).

43.	Lynch, M. *et al.* Genetic drift, selection and the evolution of the mutation rate. *Nature Reviews Genetics 2016 17:11* **17**, 704–714 (2016).

44.	Kimura, M. *The Neutral Theory of Molecular Evolution*. (Cambridge University Press, 1983). doi:10.1017/CBO9780511623486.

45.	KIMURA, M. ON THE PROBABILITY OF FIXATION OF MUTANT GENES IN A POPULATION. *Genetics* **47**, 713–719 (1962).

46.	Crow, J. and Kimura, M. An Introduction to Population Genetics Theory [Paperback]. 608 (2009).

47.	Harris, C. R. *et al.* Array programming with NumPy. *Nature 2020 585:7825* **585**, 357–362 (2020).

48.	Wickham, H. *ggplot2: Elegant Graphics for Data Analysis*. (Springer-Verlag New York, 2016).

49.	Rockel, S., Geertz, M. & Maerkl, S. J. MITOMI: A microfluidic platform for in vitro characterization of transcription factor-DNA interaction. *Methods in Molecular Biology* (2012) doi:10.1007/978-1-61779-292-2_6.

50.	Berens, C. & Hillen, W. Gene regulation by tetracyclines. Constraints of resistance regulation in bacteria shape TetR for application in eukaryotes. *Eur J Biochem* **270**, 3109–3121 (2003).

51.	Nguyen, T. N. M., Phan, Q. G., Duong, L. P., Bertrand, K. P. & Lenski, R. E. Effects of carriage and expression of the Tn10 tetracycline-resistance operon on the fitness of Escherichia coli K12. *Mol Biol Evol* **6**, 213–225 (1989).

52.	Eckert, B. & Beck, C. F. Overproduction of transposon Tn10-encoded tetracycline resistance protein results in cell death and loss of membrane potential. *J Bacteriol* **171**, 3557–3559 (1989).

53.	Rajer, F. & Sandegren, L. The Role of Antibiotic Resistance Genes in the Fitness Cost of Multiresistance Plasmids. *mBio* **13**, (2022).

54.	Poelwijk, F. J., Kiviet, D. J., Weinreich, D. M. & Tans, S. J. Empirical fitness landscapes reveal accessible evolutionary paths. *Nature* (2007) doi:10.1038/nature05451.

55.	Weinreich, D. M., Watson, R. A. & Chao, L. Perspective: Sign epistasis and genetic constraint on evolutionary trajectories. *Evolution* (2005).

56.	Greene, D. & Crona, K. The Changing Geometry of a Fitness Landscape Along an Adaptive Walk. *PLoS Comput Biol* **10**, e1003520 (2014).
